# Lineage-specific histories rather than shared drivers shape olfactory evolution in New World monkeys

**DOI:** 10.64898/2026.11.05.764729

**Authors:** J. Blumina Romero, S. Ivan Perez, Leandro Aristide

**Author notes:** Corresponding author: Leandro Aristide.

## Abstract

Variation in primate olfactory receptor gene repertoires has been linked to shared ecological and sensory drivers, including diet, diurnality, and colour vision. Yet within anthropoids, each link rests on a single lineage, leaving unclear whether this variation reflects a shared response to these drivers or lineage-specific histories. Here, we combine genomic data from 80 New World monkey species with CT-based olfactory anatomy and ecological data to investigate the evolutionary processes underlying repertoire variation. Ecological and sensory variables explain little of this variation; even routinely trichromatic, leaf-eating howler monkeys have repertoires of unremarkable size. Instead, convergent reductions in repertoire size arose through distinct genomic routes: extensive gene inactivation in titi monkeys and large structural deletions in squirrel monkeys. Owl monkeys, the only anthropoids that reverted to nocturnality, retained rather than expanded a large ancestral repertoire shared with diurnal capuchins, and instead enlarged their olfactory bulb. Repertoire size covaries with olfactory bulb volume across the radiation, but only partially within lineages: squirrel monkeys show the converse, losing receptors without a matching bulb reduction. Thus, olfactory evolution in New World monkeys reflects historical contingency rather than a shared response, with convergent changes in repertoire size and partial decoupling of genomic and anatomical evolution.

## Introduction

The evolution of olfaction in primates, particularly in anthropoids, has long been associated with a reduction in olfactory capabilities compared to other mammals (King and Fobes 1974; Baron et al. 1983). This reduction has often been linked to a shift towards visual specialisation, with olfaction progressively degenerating following the transition to diurnal lifestyles (Barton et al. 1995) and the acquisition of trichromatic vision (Gilad et al. 2004). Consistent with this sensory trade-off, anthropoids show reduced olfactory structures (T. D. Smith et al. 2007; Martinez et al. 2025; Baron et al. 1983) and smaller functional olfactory receptor (OR) gene repertoires than many other mammals (Rouquier et al. 2000; Policarpo et al. 2024; Niimura et al. 2014), with the largest reductions at the origin of haplorhines (Matsui et al. 2010; Niimura et al. 2018). OR genes encode the receptors that bind odorants, and their number is often taken as a proxy for olfactory capacity (Niimura 2012).

Beyond this reduction, however, what drives variation in repertoire size remains unclear. Once the strepsirrhine-haplorhine split is accounted for, neither activity pattern nor color vision predicts repertoire size (Niimura et al. 2018). Within anthropoids, ecological explanations rest on single lineages: leaf-eating colobines for diet (Niimura et al. 2018), the nocturnal owl monkeys for activity pattern (Niimura et al. 2018), and the routinely trichromatic howler monkeys for color vision (Gilad et al. 2004). Other changes, such as the independent acceleration of loss in each hominoid species (Niimura et al. 2018), or the deletion of an OR cluster in orangutans (Matsui et al. 2010), have no evident ecological correlates. This variation could reflect a shared response, in which lineages exposed to similar sensory and ecological conditions gain or lose receptors to a similar extent, or lineage-specific histories shaped by random events and by each lineage’s past (Langerhans and DeWitt 2004; Blount et al. 2018). Because the relevant contrasts each rest on a single lineage, cross-species associations cannot tell these alternatives apart (Uyeda et al. 2018).

Distinguishing them requires knowing not only how large repertoires are, but also how they originated, as similar outcomes can be reached by different routes (Losos 2011; Stern 2013). OR repertoires change through gene inactivation, and duplication and loss of individual genes or several clustered genes at once (Niimura and Nei 2007; Nei et al. 2008; Nozawa et al. 2007; Dong et al. 2009), while retained receptors can diversify in sequence and tuning (Chi et al. 2025). These processes leave distinct genomic traces, revealing whether a large repertoire was expanded or retained from an ancestor, and whether losses accumulated progressively or through a few large deletions. Under a shared response, gains and losses should track the conditions each lineage experienced. Under lineage-specific histories, repertoires would instead diverge independently of these conditions, for example, when a few large deletions, rather than many inactivations, make the extent of loss depend on rare events.

A shared response also predicts how olfactory anatomy should change. OR repertoire size covaries with olfactory bulb volume and turbinal number across mammals and primates (Martinez et al. 2025; Christmas et al. 2023; Niimura et al. 2018). If repertoire changes because reliance on olfaction changes, olfactory anatomy should change along the same branches. Instead, if repertoires change without a change in reliance, or the two components respond to different selective pressures and constraints (Niimura 2009; Yohe and Krell 2023), they may change independently within lineages while still covarying across species, a distinction with implications for inferring olfactory capacity from bulb endocasts in fossil species (Martinez et al. 2025; Lang et al. 2026).

New World monkeys (platyrrhines), which originated ∼25 Ma and diversified across Central and South America into a wide range of ecological niches (Perez et al. 2013; Kay 2015; A. L. Rosenberger 1992), allow these alternatives to be examined within a single radiation. They include two lineages central to ecological explanations: *Aotus*, the only nocturnal/cathemeral anthropoid genus, whose reversal from a diurnal ancestor involved extensive remodelling of its visual system (Ankel-Simons and Rasmussen 2008), and *Alouatta*, the only routinely trichromatic and the most folivorous platyrrhine (Gilad et al. 2004; A. L. Rosenberger 1992). Under a shared response, *Aotus* should have gained receptors, or lost fewer than its diurnal relatives, after becoming nocturnal, and *Alouatta* should have lost more than other platyrrhines. More generally, the largest and smallest repertoires should correspond to ecological or sensory contrasts.

Here, we investigate the evolutionary history of OR repertoires across 80 platyrrhine species, representing all families and nearly all genera, combining phylogenetic comparative genomics with CT-based cranial anatomy and ecological data. The large repertoire of the nocturnal *Aotus* was not gained with nocturnality but retained from the platyrrhine ancestor, as in the diurnal capuchins. The smallest repertoires arose through distinct routes, extensive pseudogenization in titi monkeys and large structural deletions in *Saimiri*, the latter despite a generalist ecology shared with capuchins. Repertoire size and olfactory bulb volume covary across the radiation but partially decouple in both *Aotus*, whose bulb enlarged without new receptors, and *Saimiri*, which lost receptors without a matching reduction of the bulb. *Alouatta*, despite routine trichromacy and folivory, has a repertoire of unremarkable size, and positive selection on the remaining receptors varies among lineages largely independently of repertoire change. Together, these findings show that, although repertoires changed mainly through loss from a large ancestral repertoire, olfactory evolution in platyrrhines reflects the outcome of historically contingent changes within individual lineages rather than a shared response to ecological or sensory drivers.

## Results

### OR repertoire size and composition are conserved within genera and divergent among a few lineages

To characterize variation in the platyrrhine OR gene repertoire, we first identified and characterized OR loci across the genomes of 80 species (Supplementary Table 1). Functional repertoire size (corrected intact ORs; see Methods) ranged from 322 to 450 genes and pseudogene number ranged from 89 to 219 (Fig. 1a). Both counts were strongly phylogenetically structured and highly conserved within genera. The largest functional repertoires were found in *Aotus* and capuchins (*Cebus* and *Sapajus)*, which had remarkably similar high numbers of functional ORs (425-447 and 414-450, respectively), although *Aotus* carried fewer pseudogenes than capuchins (94-115 versus 138-162). In contrast, *Saimiri*, the sister lineage to the capuchins, had the smallest functional repertoires (322-340) together with reduced pseudogene counts (89-141). This contrast is particularly striking because *Saimiri* and capuchins are closely related and share a broadly generalist feeding ecology (A. L. Rosenberger 1992). The callicebins (titi monkeys; *Plecturocebus* and *Cheracebus*) had similarly small functional repertoires (329-354), but the highest pseudogene numbers in the radiation (159-219).

**Figure 1.**
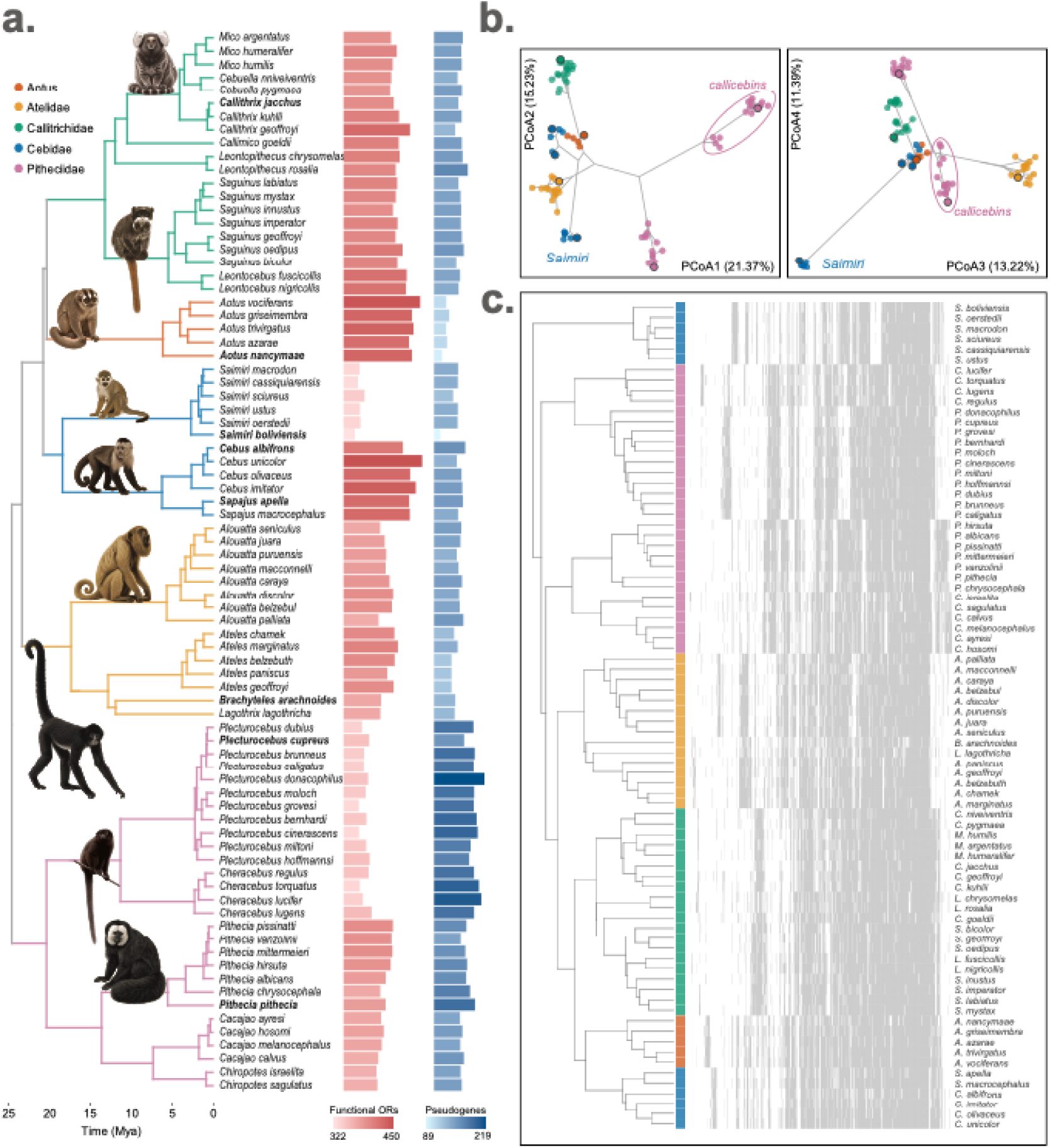
Olfactory receptor (OR) repertoire size and composition across 80 platyrrhine species. (**a)** Time-calibrated phylogeny of the 80 species, colored by family, with the number of functional OR genes (red bars) and OR pseudogenes (blue bars) in each species. Counts are corrected for assembly fragmentation (see Methods). Raw and corrected intact, truncated and pseudogenized gene counts for all species are provided in Supplementary Data 1. Species in bold have high-quality assemblies (Contig N50 > 10 Mb). Illustrations depict representative genera. **(b)** Principal coordinate analyses (PCoA) of Jaccard dissimilarities in orthogroup presence-absence among species, based on the 738 orthogroups containing at least one intact OR (pseudogenes excluded). Left, PCo1 against PCo2; right PCo3 against PCo4; percentages in axis labels indicate the variance explained by each axis. Points are species colored by family, with high-quality assemblies outlined in black; grey lines show the phylogenetic relationships projected on the compositional space. **(c)** Presence (grey) or absence (white) of each orthogroup (columns) in each species (rows), with species ordered by UPGMA clustering of the same dissimilarities (dendrogram, left).

Because similar gene counts may mask differences in composition, which may reflect functional and evolutionary differences, we next assigned OR sequences from all 80 species to orthogroups: sets of genes descending from a single ancestral gene in a set of species. The analysis grouped platyrrhine ORs into 738 orthogroups with at least one intact gene, of which 345 were shared across all five platyrrhine families. We compared orthogroup presence-absence among species using Jaccard dissimilarities, principal coordinate analysis (PCoA) and hierarchical clustering. Repertoire composition was strongly phylogenetically structured (Fig. 1b,c), almost recreating the species tree (Fig. 1a), with species within the same genus occupying tight regions of the compositional space (Fig. 1b,c). This also indicates that the analysis is robust to differences in genome assembly quality. Notably, *Saimiri* was not only separated from the capuchins, but also showed a divergent repertoire from all other platyrrhines (Fig. 1b,c), while its sister clade, the capuchins, clustered together with *Aotus* (Fig. 1b,c), despite their contrasting activity patterns and sensory ecologies. Within Pitheciids, the titis differed markedly from the pithecines, and also from the similarly small-sized repertoires of *Saimiri*.

Size and composition thus converge on the same picture: both are conserved within genera and largely track phylogenetic history, but a few lineages break this structure. These lineages carry the largest and the smallest repertoires in the radiation, and depart from phylogenetic expectations either by diverging sharply from their closest relatives, or by resembling distantly related lineages.

### Distinct genomic mechanisms underlie OR repertoire evolution in platyrrhines

#### Limited role of pseudogenization in repertoire contraction

If repertoire contraction proceeded mainly through gene inactivation (pseudogenization), species with fewer functional ORs should carry more pseudogenes, although a perfect relationship is not expected as pseudogenes can also suffer duplication and loss (Matsui et al. 2010). Across platyrrhines, pseudogene number was weakly and negatively associated with functional OR number (PGLS β = −0.65, *p* = 0.043; Pagel’s λ = 0.91; R² = 0.05; Fig. 2a). This is consistent with pseudogenization contributing to repertoire contraction, but accounts for little of the variation in repertoire size across the radiation, as the smallest repertoires illustrate. The titis, despite having functional repertoires similar in size to those of *Saimiri*, carry the highest pseudogene counts in the radiation (Figs. 1a, 2a), as expected if their contraction proceeded largely through pseudogenization. *Saimiri* shows the opposite pattern. Adding a *Saimiri* term to the model improved its fit (LRT p = 0.027; Supplementary Table 3): the genus carried ∼29% fewer pseudogenes than expected for its functional repertoire size (β = −0.34, *p* = 0.024), and once this was accounted for, the relationship between the two counts also became steeper (β = −0.95, p = 0.01; R^2^ = 0.10). The effect was in the same direction when the analysis was restricted to the high-quality assemblies (β = −0.77), although statistical power was limited at this sample size (Supplementary Table 3). Thus, the smallest functional repertoires in the radiation were reached by different routes: pseudogenization can account for the titi pattern, whereas the reduced repertoire of *Saimiri* is not fully explained by gene inactivation, suggesting that additional mechanisms of loss were involved.

**Figure 2.**
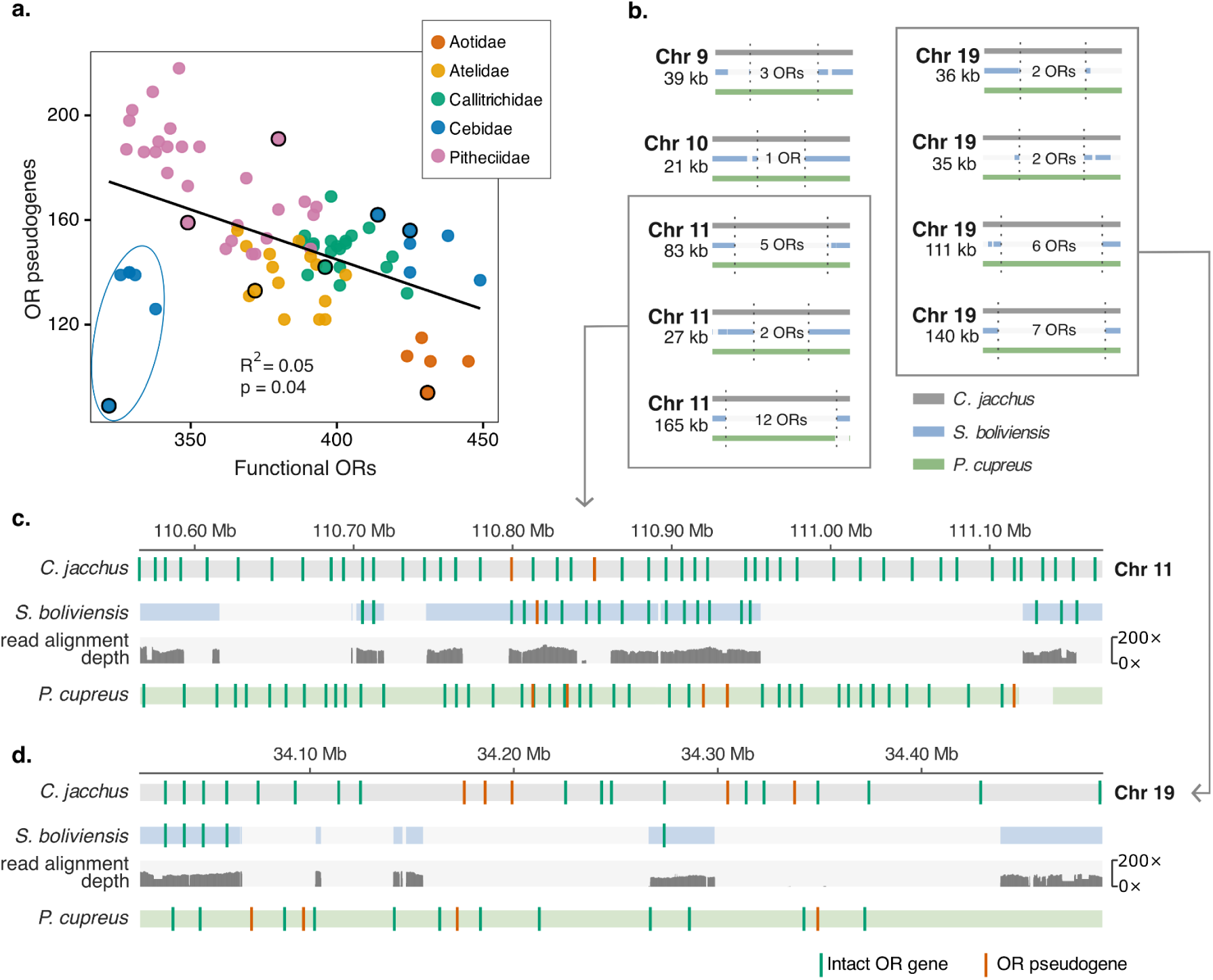
Functional and pseudogene OR counts across platyrrhines, and OR-containing regions absent from the *Saimiri boliviensis* genome. (**a)** Number of OR pseudogenes against number of functional ORs for the 80 species (counts corrected for assembly fragmentation), coloured by family. The black line shows the phylogenetic generalized least-squares (PGLS) fit, estimated on log-transformed counts and plotted in original units. *Saimiri* species are circled in blue, and species with high-quality assemblies are outlined in black. **(b)** The nine OR-containing regions of the *Callithrix jacchus* genome that lack aligned sequence in *S. boliviensis* but are aligned in *Plecturocebus cupreus*. Each region is shown as the reference *C. jacchus* (grey), and aligned *S. boliviensis* (blue) and *P. cupreus* (green), with its *C. jacchus* chromosome, its length and the number of ORs it contains. Boxes group the regions on chromosomes 11 and 19 shown in detail in c and d. **(c, d)** Detail of the regions on *C. jacchus* chromosomes 11 (c) and 19 (d). Tracks show, from top, the *C. jacchus* reference, *S. boliviensis* segments aligned to it, the depth of *S. boliviensis* PacBio HiFi reads mapped to the *C. jacchus* reference (0–200×), and *P. cupreus* segments aligned to it. Vertical lines mark annotated ORs (green, intact; red, pseudogenes). Coordinates refer to the *C. jacchus* assembly.

#### Large structural deletions as an alternative route to repertoire contraction

A small functional repertoire accompanied by few pseudogenes suggests that OR loci were removed rather than inactivated. Because OR genes are organized in genomic clusters (Niimura 2012), we hypothesized that large structural deletions encompassing OR-containing regions underlie the repertoire contraction in *Saimiri*. To test this, we searched for OR-containing regions of the *Callithrix jacchus* genome lacking aligned sequence in the long-read assembly of *Saimiri boliviensis*, using *Plecturocebus cupreus* as an outgroup to distinguish losses in *Saimiri* from gains in *Callithrix*. We initially identified 22 candidate OR-containing regions in the *C. jacchus* genome absent in *S. boliviensis*. We then mapped raw *S. boliviensis* reads to the *C. jacchus* assembly and found that 15 of such regions had a near-complete drop in read alignment depth relative to their 20 kb flanks (gap-to-flank depth ratio < 0.1), strongly suggesting the absence of the underlying sequence in *Saimiri* (Fig. 2b,c). Furthermore, 9 of the 15 regions aligned over more than 80% of their length to the *P. cupreus* genome, supporting their interpretation as *Saimiri*-specific losses of ancestral sequence (Supplementary Table 4). These 9 regions contained 34 functional ORs and 6 pseudogenes in *C. jacchus,* whereas the corresponding regions in *P. cupreus* contained 30 functional ORs and 6 pseudogenes. Notably, 7 of the 9 regions fell within two major *C. jacchus* OR clusters (Fig. 2b). Deleted sequence stretches ranged from ∼21-165 kb, indicating the loss of large genomic segments rather than isolated loci (Fig. 2c). Orthology analyses further supported gene loss rather than relocation: 30 out of 34 functional ORs within these *C. jacchus* regions lacked an identifiable ortholog elsewhere in the *S. boliviensis* genome (Supplementary Table 5), representing approximately 41% of the difference in functional repertoire size between the two species. Together, these results identify large structural deletions of OR-containing regions as a major contributor to repertoire contraction in *Saimiri*.

#### Retention rather than expansion underlies the largest repertoires

The large repertoires of *Aotus* and the capuchins could reflect either extensive retention of ancestral OR diversity or independent lineage-specific expansions. Their compositional similarity (Fig. 1b,c) suggested shared retention, which we tested by estimating how much of the ancestral platyrrhine repertoire each lineage retains. Using Dollo’s parsimony on orthogroup presence/absence, we inferred that the platyrrhine ancestor had 519 orthogroups containing at least one intact OR. Among the eight species with high-quality assemblies, retention was highest in *Aotus nancymaae* (73.4%), *Sapajus apella* (71.9%), and *Cebus albifrons* (68.2%) (Fig. 3; Supplementary Table 6). *S. boliviensis* retained the fewest (56.1%) and lost the most, with 188 ancestral orthogroups (36.2%) absent either as intact genes or pseudogenes, consistent with the deletions identified above.

**Figure 3.**
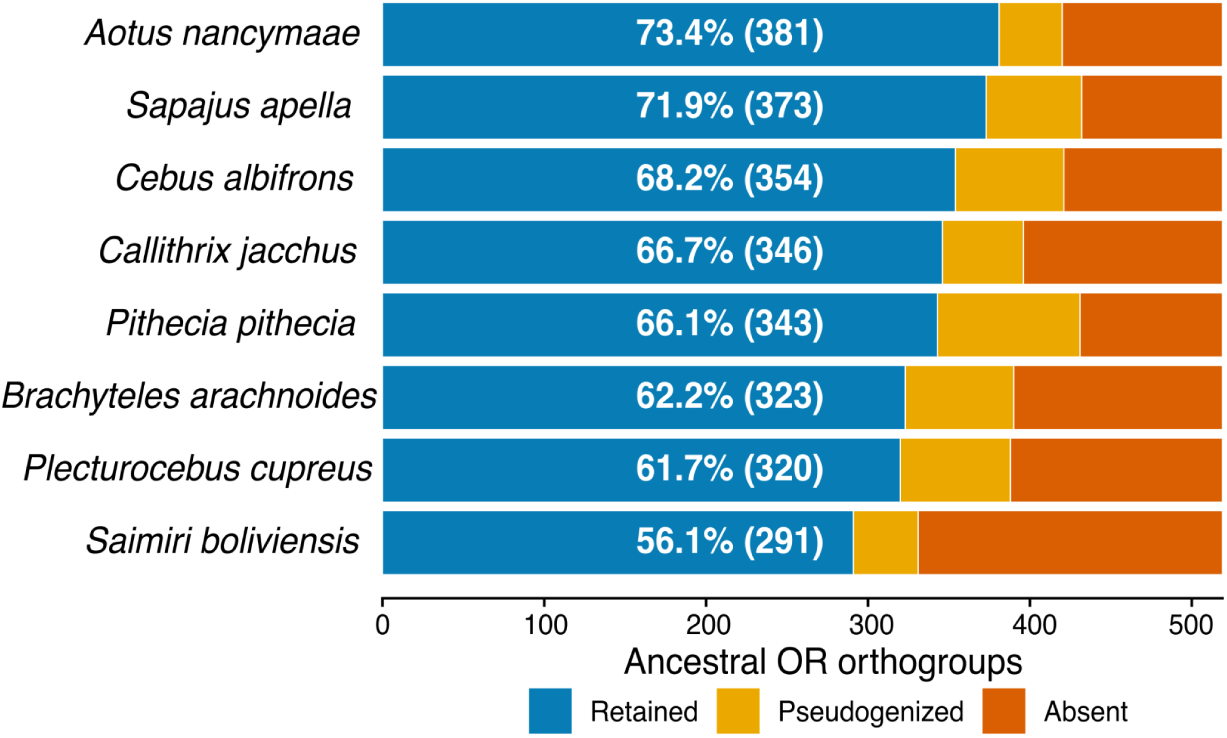
Retention of the ancestral platyrrhine OR repertoire in species with high-quality genome assemblies. For each of the eight species with high-quality assemblies, the bar divides the 519 orthogroups inferred to be present in the platyrrhine ancestor (Dollo parsimony across the 80 species; see Methods) into those retained with at least one intact OR (blue), those present only as pseudogenes (yellow) and those absent (orange). Labels give the number and percentage of ancestral orthogroups retained in each species.

To test the alternative explanation, we built a phylogeny of all functional OR sequences from six selected species with high-quality assemblies and searched for species-specific monophyletic clusters in *Aotus* or capuchins. We found no evidence of large recent expansions: the largest supported (bootstrap ≥ 95%) lineage-specific clusters contained three ORs in capuchins (shared by *C. albifrons* and *S. apella*) and two in *A. nancymaae* (Supplementary Table 7). Thus, recent duplications occurred in both lineages but were too limited to account for their large repertoires. Together, high retention of ancestral orthogroups and the absence of large duplications indicate that the large OR repertoires of *Aotus* and the capuchin monkeys primarily reflect a retention of the ancestral repertoire rather than parallel expansion.

#### Sequence-level positive selection varies among lineages independently of repertoire change

Beyond variation in repertoire size and composition, OR diversification can also proceed through changes in the coding sequences of functional receptors, potentially modifying odorant binding and recognition (Chi et al. 2025). We therefore characterized selection on functional OR sequences across the radiation, analyzing each orthogroup separately in one species per genus (20 species). Under the CODEML free-ratio model, most terminal branches had ω < 1 (84.8% of 7,540 valid estimates, and 92.5% in seven high-quality genomes with comparable branch lengths; Fig. 4a; Supplementary Data 3; Supplementary Fig. 5), consistent with predominantly purifying selection in all lineages.

**Figure 4.**
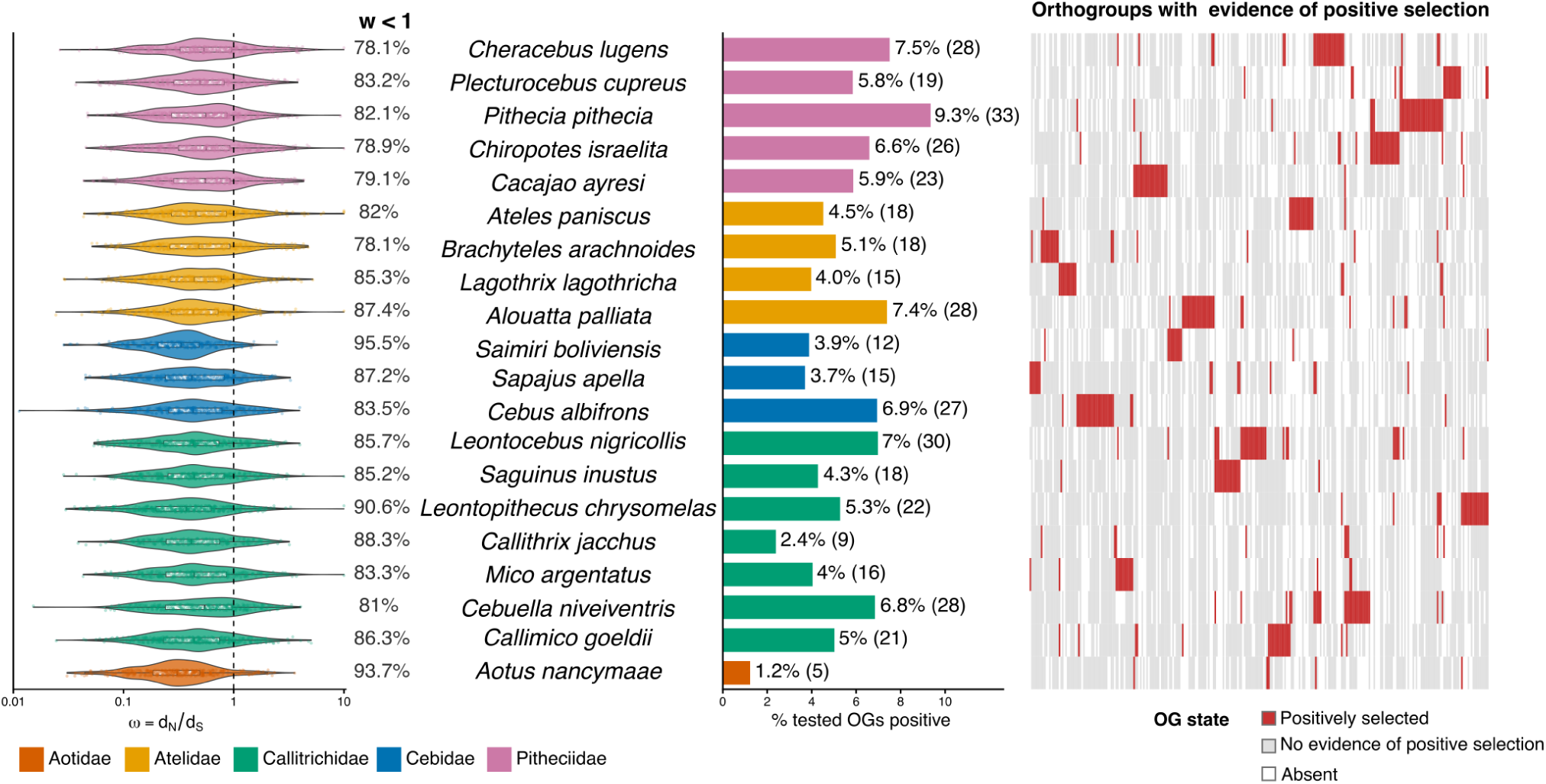
Selection on OR coding sequences across 20 platyrrhine genera. Species (centre), one per genus, are grouped by family; colours indicate families as in Fig. 1. **(a)** Distribution of terminal-branch ω (dN/dS) estimated for each species with the CODEML free-ratio model (log scale); the dashed line indicates ω = 1. Percentages give the proportion of terminal branches with ω < 1 in each species, including estimates at the lower bound of estimation, which are not plotted; branches for which ω could not be estimated are excluded (see Methods). **(b)** Percentage of tested orthogroups with at least one terminal branch under episodic positive selection in each species (aBSREL; BH-adjusted p < 0.05), with the numbers of such orthogroups in parentheses. **(c)** aBSREL results for the 283 orthogroups with at least one significant terminal branch, with species as rows, in the same order as in a and b, and orthogroups as columns. Red, significant result; grey, orthogroup present without a significant result; white, orthogroup absent from the species.

Against this constrained background, aBSREL detected episodic positive selection on 417 of 8,362 tested terminal branches (4.99%; BH-adjusted p < 0.05; Supplementary Data 4), distributed across 283 of 603 orthogroups analysed (46.9%). Positive selection was dispersed across the repertoire: 192 of the 283 positive orthogroups (67.8%) contained a single significant branch, and only 30 had significant branches in three or more species (Fig. 4c), suggesting that sequence-level diversification has involved different ORs in different species rather than repeatedly affecting a common subset. The results were robust to removal of truncated ORs (Supplementary Fig. 6).

The frequency of positive selection, however, differed among lineages, from 1.2% of tested orthogroups in *A. nancymaae* to 9.3% in *Pithecia pithecia* (Fig. 4b). All of the five pitheciid genera fell in the upper part of this range (5.8–9.3%), while the remaining genera spanned nearly all of it (1.2-7.4%). The high frequencies also extended to the titis, which do not share the seed predation specialization of pitheciines, indicating that this pattern is not tied to a single dietary specialization. The same ranking held in the seven-species analysis (Supplementary Fig. 5).

Lineages with similar repertoires can differ sharply in sequence-level selection. *A. nancymaae* and *C. albifrons*, which retain among the largest fraction of the ancestral repertoire (Fig. 3), differed about five-fold in the frequency of positive selection (1.2% vs. 6.9%; 1.6% vs. 8.2% in the seven-species analysis). For *Aotus*, the reversal to nocturnality was thus accompanied neither by extensive change in OR gene content nor by elevated positive selection on OR sequences. Together, these results indicate that sequence-level adaptation represents a distinct layer of OR evolution, varying among lineages largely independently of the processes that shape repertoire size and composition.

### OR repertoire size and olfactory anatomy covary across the radiation but can decouple within lineages

To ask whether the repertoire histories identified above are reflected in olfactory anatomy, we characterized olfactory morphology from µCT scans of 59 individuals representing

33 platyrrhine species (Supplementary Data 5), measuring olfactory bulb volume (OBV), cribriform plate area (CPA), turbinal surface area (TSA) and olfactory turbinal number. We examined the relationship between genomic and anatomical components at two phylogenetic scales: whether functional OR repertoire size covaries with olfactory anatomy across the radiation, and whether lineages with distinct repertoire histories follow that shared relationship. Two lineages allow the second question to be posed from opposite directions. In *Saimiri*, the repertoire contracted through structural deletions. In *Aotus*, the ancestral repertoire was retained through the reversal to nocturnality, but the olfactory bulbs are the largest relative to endocranial volume of any living anthropoid, a condition interpreted as derived within the otherwise diurnal radiation (Lang et al. 2026). Capuchins, which retain a similar ancestral repertoire but are diurnal, provide a comparison for *Aotus*.

Across platyrrhines, with body mass (BM) and endocranial volume (ECV) as allometric covariates, OBV was the trait most strongly associated with repertoire size (β = 0.19, R² = 0.52, p < 0.01). CPA and turbinal number showed weaker associations (β = 0.06, R² = 0.16, p = 0.04; β = 0.07, R² = 0.25, p < 0.01), and TSA showed none (β = 0.02, R² = 0.04, p = 0.38; Supplementary Table 8). We therefore examine lineage-specific departures in bulb volume with three models (Fig. 5a-c). Relative to BM and ECV, *Aotus* had larger bulbs than expected (β = 0.70, p < 0.01), whereas *Saimiri* had somewhat smaller bulbs (β = -0.51, p = 0.06; Fig. 5a) and capuchins had bulbs of unremarkable size (Fig. 5a). Adding functional OR number as a predictor (Supplementary Table 8) did not account for the larger bulbs of *Aotus*, which remained ∼43% larger than expected (β = 0.36, p < 0.01; Fig. 5b). In *Saimiri*, the deviation changed sign: its bulbs were reduced less than its small repertoire predicted and were, if anything, larger than expected for that repertoire (∼32%; β = 0.28, p = 0.05).

**Figure 5.**
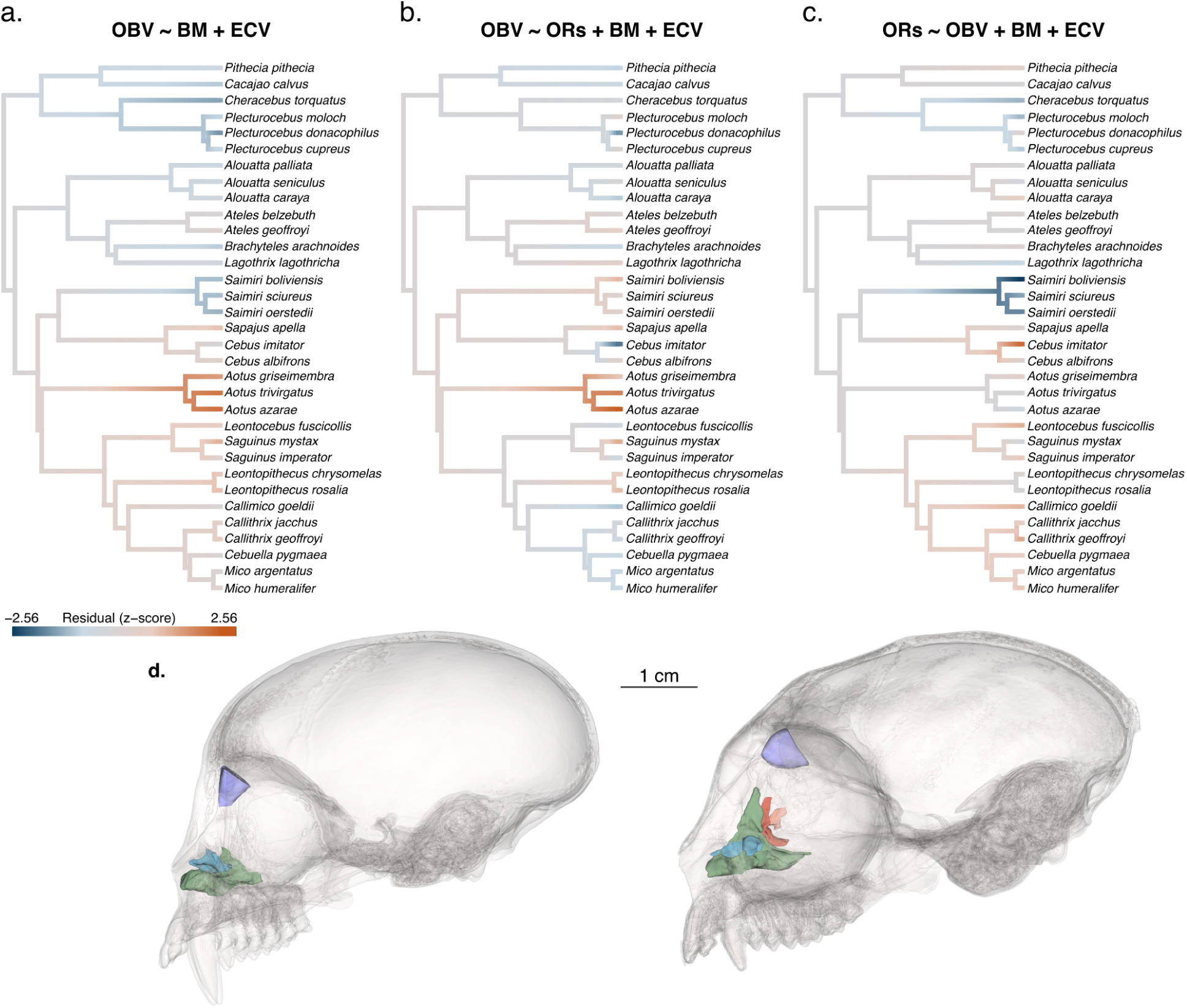
Evolution of olfactory bulb volume and OR repertoire size across platyrrhines. (**a-c**) Standardized residuals (z-scores; blue, negative; red, positive) from three phylogenetic generalized least-squares (PGLS) models relating olfactory bulb volume (OBV), number of functional ORs (ORs), body mass (BM) and endocranial volume (ECV) across the 33 species with morphological data. Residuals are from models without genus terms; tip colors show the residuals and branch colors their reconstruction under Brownian motion. **(a)** OBV relative to BM and ECV (OBV ∼ BM + ECV). **(b)** OBV relative to OR number, BM and ECV (OBV ∼ ORs + BM + ECV). **(c)** OR number relative to OBV, BM and ECV (ORs ∼ OBV + BM + ECV). **(d)** Three-dimensional reconstructions of the crania of *Saimiri boliviensis* (left) and *Aotus azarae* (right) in lateral view, showing the olfactory bulb endocast (purple), the left nasoturbinal (blue) and the left ethmoturbinals (green, red and pink). Scale bar, 1 cm.

Both lineages therefore have bulbs larger than expected for their repertoire, a pattern that can arise from an enlarged bulb or from a reduced repertoire. The two can be distinguished by reversing the model: because repertoire size and bulb volume are only partially correlated, a lineage deviates more clearly, relative to the residual scatter, when the trait that departed is the response variable. With repertoire size as the response, *Aotus* had the repertoire expected from its bulb (β = -0.01, p = 0.91), whereas *Saimiri* had ∼12% fewer functional ORs than predicted from its bulb (β = -0.13, p < 0.01; Fig. 5c) and capuchins somewhat more than predicted (β = 0.09, p = 0.04; Fig. 5c).

Combined with the repertoire histories reconstructed above, the two directions of the analysis therefore indicate which trait changed in each lineage. In *Saimiri*, the departure lies in the repertoire, which lost OR loci through structural deletions and was reduced more than the bulb. In *Aotus*, it lies in the bulb: *Aotus* and the capuchins carry similar repertoires retained from the platyrrhine ancestor (Figs. 1b, 3), but only *Aotus* has an enlarged bulb. Thus, although repertoire size and bulb volume covary across platyrrhines, they are partially decoupled in *Saimiri* and *Aotus*, with a different component departing in each.

### Ecological and sensory variables explain little repertoire variation

Finally, we asked whether repertoire variation across the radiation also reflects a shared response to ecological or sensory drivers, as proposed for primates and mammals. Despite the considerable dietary diversity of platyrrhines, even among congeneric species (Supplementary Fig. 4; Supplementary Data 2), neither functional nor pseudogene counts were associated with the proportion of fruits, seeds, leaves, exudates, or animal prey in the diet, including in models adding activity pattern or colour vision (BH-adjusted p ≥ 0.75 for all PGLS models; Supplementary Table 2). In contrast, activity pattern and visual phenotype provided the only detectable associations: diurnal species had fewer functional ORs and more pseudogenes than nocturnal/cathemeral species, and higher color vision scores were associated with more pseudogenes, though not with functional OR counts. All these models explained little variation, however (R² = 0.06-0.22; Supplementary Table 2), and the associations rest on a single evolutionary transition, as *Aotus* species are the only nocturnal/cathemeral and monochromatic platyrrhines. Ecology and phylogeny are strongly collinear in platyrrhines (Alfred L. Rosenberger et al. 2008), so ecological effects are difficult to distinguish from similarity due to shared ancestry. Because these models estimate a single relationship across the radiation, weak or absent associations argue against a common ecological effect, not against effects confined to particular lineages. The variation in OR repertoire size observed across platyrrhines, however, does not correspond to contrasts in any predictor examined: *Aotus* shares its retained repertoire with the diurnal capuchins, and *Saimiri* contracted its repertoire despite sharing their generalist diet and diurnal activity. They are better explained by the lineage-specific histories reconstructed above than by shared ecological or sensory drivers. Conversely, *Alouatta*, which combines routine trichromacy with the most folivorous diet in the radiation, shows no departure: its functional repertoire (367–395 ORs) lies near the radiation median (390; Fig. 1a). Its pseudogene fraction (26-30%), the measure on which the link to trichromacy was originally based (Gilad et al. 2004), is slightly above that of other atelids (23-26%) but typical of platyrrhines with polymorphic color vision (median 28%).

## Discussion

By evaluating primate olfaction within a single radiation, our results allowed us to better understand the evolution of OR repertoires. Platyrrhine repertoires were shaped by distinct histories of gene loss rather than by a shared response to ecological or sensory drivers: loss is common to all lineages, but its extent, mechanism and consequences for the rest of the olfactory system are lineage-specific. Three general patterns emerge from these results. First, gene counts alone do not identify the processes that produced them: similar repertoire sizes were reached by distinct routes, and the largest repertoires reflect shared retention rather than expansion. Second, the association between repertoire size and olfactory bulb volume that holds across the radiation was only partially maintained within lineages, and in opposite directions. Third, ecological and sensory variables accounted for little of this variation, and diet for none of it.

The first of these patterns is clearest in the two smallest repertoires. Titi repertoires contracted through extensive pseudogenization, the predominant route of OR loss in mammals (Niimura and Nei 2007; Nei et al. 2008; Hughes et al. 2018). They also showed some of the most frequent positive selection in the radiation: the inactivation of many receptors has been accompanied by the diversification of others, a pattern suggesting reorganization rather than pure decline. *Saimiri* reached similar-sized functional repertoires with fewer pseudogenes, less frequent positive selection and a distinct composition, in large part through deletions of OR-containing regions spanning tens to hundreds of kilobases. This adds OR evolution to a growing body of evidence that structural variants are a major source of genomic change in primates (Mao et al. 2024), and other organisms (Mérot et al. 2020; Gompert et al. 2025). As in orangutans, which lost a large OR cluster (Matsui et al. 2010), much of the *Saimiri* contraction suggests a few events of large effect rather than many independent inactivations. The convergence of the two smallest repertoires is thus confined to gene counts, an instance of similar phenotypes arising through distinct genetic routes (Losos 2011; Stern 2013). Comparisons with the capuchins show that the largest and smallest repertoires do not correspond to ecological or sensory contrasts: *Saimiri* shares their generalist ecology but not their repertoire, whereas *Aotus* shares their repertoire despite a different activity pattern and color vision, through retention from the platyrrhine ancestor rather than convergent expansion. Similar repertoire sizes can therefore be convergent or inherited, and ecologically similar lineages can diverge through lineage-specific events such as the deletions in *Saimiri* (Langerhans and DeWitt 2004; Blount et al. 2018).

The second pattern emerges from one of the most reproducible results in olfactory evolution, the phylogenetic association between receptor number and relative olfactory bulb size (Martinez et al. 2025; Policarpo et al. 2024; Niimura et al. 2018; Chi et al. 2025), which we recovered within platyrrhines, along with the association with turbinal number found across placental mammals (Christmas et al. 2023). Turbinal surface area showed none, consistent with the heterogeneous distribution of olfactory epithelium in the primate nasal cavity (T. D. Smith et al. 2004). Each olfactory sensory neuron typically expresses a single OR, and neurons expressing the same receptor project to a few glomeruli in the olfactory bulb (Tufo et al. 2022; Mombaerts et al. 1996). However, glomerular number is not fixed by receptor number, as *Callithrix jacchus* has a mouse-like glomerular count despite a much smaller repertoire (Moriya-Ito et al. 2015). Bulb volume therefore depends not only on the number of receptor types but also on how many sensory neurons and glomeruli correspond to each, whereas repertoire size counts receptor types alone. How either maps onto olfactory performance remains unresolved, as neither reliably predicts sensitivity across primates (Laska and Salazar 2015), but sensitivity has been linked to the number of sensory neurons and the pooling of their input onto glomeruli (Martinez et al. 2025). The two would covary across the radiation if lineages that rely less on olfaction tend both to lose receptors and to reduce their bulbs, and would separate within lineages when only one of them changes.

Within lineages, *Saimiri* and *Aotus* show the two directions in which the components can separate. In *Saimiri*, the repertoire contracted while the bulb was reduced only partially, in a generalist lineage that retains high olfactory sensitivity (Laska et al. 2000; Laska et al. 2005), consistent with largely unchanged reliance on olfaction. Because this offset is shared by all sampled species and absent from the capuchins, the contraction predates the divergence of the extant species, making a transient lag between genomic and anatomical change unlikely. In *Aotus*, the olfactory bulb, the largest relative to endocranial volume of any living anthropoid (Lang et al. 2026), is larger than its repertoire predicts and than those of the capuchins, which carry an equivalent repertoire. The enlargement is likely derived, as the early Miocene *Tremacebus harringtoni*, a proposed relative of *Aotus*, had a smaller bulb (Kay et al. 2004). The reversal to nocturnality therefore involved the olfactory system, but through its bulb rather than its receptors. Chi et al. (2025) linked narrowly tuned receptors to dim-light activity, so a return to nocturnality might have been expected to re-elaborate them. Instead, *Aotus* retains more of the ancestral platyrrhine repertoire, which Chi et al. inferred to be enriched in broadly tuned receptors, and shows the least positive selection in the radiation. The olfactory–visual coevolution described across primates thus holds in *Aotus* for vision, where Chi et al. found dim-light-adapted rhodopsin kinetics, and for the bulb, but not for the receptors. In the Nile rat *Arvicanthis niloticus*, a shift to diurnality was followed at both levels, with reductions in nasal and bulb anatomy, OR gene loss and relaxed purifying selection (Liao et al. 2024). This suggests an asymmetry in the constraints on the two components: reduced reliance on olfaction can be readily followed by gene loss, whereas increased reliance may be met by anatomy sooner than by the gain of new receptors, which depends on duplication and divergence.

The third pattern concerns the drivers themselves. Neither visual specialization nor diet produced a shared response at this scale. The associations with activity pattern and colour vision rest on a single transition, as *Aotus* is the only nocturnal and monochromatic platyrrhine. Nocturnality has long been thought to favour reliance on olfaction, and Niimura et al. (2018) noted the larger repertoire of *Aotus* relative to *Callithrix* and *Saimiri*, but including the diurnal capuchins removes the contrast: the large repertoire is the ancestral condition, which nocturnality may have helped maintain but is not needed to explain. The question is therefore not why *Aotus* kept a large repertoire, but why most other lineages lost part of theirs. Gene loss is thus the main source of variation in repertoire size, as suggested across primates (Matsui et al. 2010; Niimura et al. 2018), but, as among hominoids, its extent differs among lineages with the same activity pattern and color vision. *Alouatta*, the only routinely trichromatic and the most folivorous platyrrhine, provides the comparison with other New World monkeys: neither its functional repertoire nor its pseudogene fraction, the basis of the link to trichromacy (Gilad et al. 2004), is unusual among platyrrhines. Links between repertoire size and folivory or herbivory (Niimura et al. 2018; Hughes et al. 2018) instead rest on lineages with no platyrrhine counterpart, most clearly colobines. Positive selection was not organized by diet either, being elevated in all pitheciids, including titis, which lack the seed predation of pitheciines (A. L. Rosenberger 1992; Norconk 2021). Dietary categories describe what a species eats rather than how much it relies on smell to find it, as even among sympatric fruit-eating platyrrhines sniffing frequency differs (Melin et al. 2022). Finer measures of sensory ecology, such as foraging mode, the reliability of visual cues and odorant chemistry (Nevo and Heymann 2015) may be more informative.

Three limitations bound these conclusions. Our sample combines assemblies of heterogeneous quality; we modelled truncation explicitly and restricted mechanism-level analyses to high-quality genomes, and the tight clustering of congeners indicates that the main patterns are not assembly artefacts, although absolute counts in fragmented assemblies remain approximate. The deletion analysis rests on the only *Saimiri* genome with long reads, so whether all *Saimiri* species carry these deletions remains to be tested. Finally, intact ORs are not necessarily expressed and pseudogenes occasionally are, so repertoire size is an indirect proxy for perceptual capacity, as endocast volume is for the bulb itself. Glomerular counts in *Aotus* and the capuchins, which share a repertoire but differ in bulb size, would show how the bulb enlarged without new receptors.

Dense sampling within a radiation can resolve what comparisons across distant taxa tend to average over and separate singular events from shared responses, providing a “phylogenetic natural history” (Uyeda et al. 2018). Components conventionally treated as coupled can change partly independently within a lineage while covarying across the radiation, so associations estimated across species describe a shared tendency rather than the history of any one lineage. This has direct implications for the use of olfactory bulb endocasts to infer olfactory capacity in fossil taxa (Lang et al. 2026; Martinez et al. 2025): the bulb tracks the receptor repertoire across lineages, but within any one of them it need not. Repertoire size, in this view, is not a position on a scale of sensory trade-offs or uniform reduction, but the outcome of historically contingent, lineage-specific changes that gene counts alone cannot recover.

## Methods

### Phylogeny

All analyses requiring a species tree used a single dated, ultrametric composite phylogeny. The backbone was obtained from the primate chronogram of Kuderna et al. (2023), pruned to platyrrhines. Six species absent from that phylogeny were added as tips following published relationships and divergence-time estimates: *Aotus nancymaae*, *Plecturocebus donacophilus*, *Pithecia pithecia*, *Saimiri boliviensis* and *Brachyteles arachnoides* using the 372-species dated tree of dos Reis *et al*. (2018) and *Cebus imitator* following Martins-Junior *et al*. (2018). This tree was pruned to the taxa available for each analysis group (80 species for genomic analyses, 51 for ecological models and 33 for morphological models).

### Genomic data

The genomic dataset comprises 80 whole-genome assemblies, one per species, representing 20 out of 21 broadly recognized platyrrhine genera (Mammal Diversity Database V 2.5, 2026) and all five families (Fig. 1). The only exception, *Callicebus*, is a recent taxonomic split within titi monkeys (Byrne et al. 2016), which are represented here by *Plecturocebus* and *Cheracebus*. All were downloaded from the NCBI Genome database; accession numbers and assembly statistics are provided in Supplementary Table 1. Seventy-two assemblies are based on Illumina short-reads, most generated at ∼35x coverage and assembled under the same pipeline as part of a large-scale primate genomic survey (Kuderna et al. 2023). The remaining eight incorporate long-read data (PacBio or Oxford Nanopore) and are substantially more contiguous; this subset includes at least one representative of each platyrrhine family and served as the anchor for all analyses. We handled the heterogeneity in assembly contiguity through sequence curation during olfactory receptor annotation and through an explicit correction of repertoire counts for truncation at contig boundaries, both described below, and we repeated the main analyses on the subset of highest-quality assemblies throughout.

### Identification and classification of OR genes

We identified and classified olfactory receptor (OR) loci in each genome assembly through similarity-based annotation, phylogenetic filtering, fragment merging and structural classification. The phylogenetic filtering step follows Policarpo et al. (2024) and uses their curated reference alignment of vertebrate ORs directly; the remaining steps were developed for this dataset, drawing on their treatment of transmembrane topology and of sequences lying close to assembly boundaries. The complete pipeline is summarized in Supplementary Fig. 1.

Candidate OR loci were annotated with BITACORA v.1.4 (Vizueta et al. 2020), which combines BLAST and profile hidden Markov model (HMM) searches and is designed to annotate and curate gene families in genome assemblies of variable quality. BITACORA was run in “genome mode” for each species assembly, using a custom reference database of OR protein sequences together with a profile HMM derived from it. The database consisted of 409 reviewed human OR sequences and 229 unreviewed platyrrhine OR sequences, retrieved from UniProt through the InterPro olfactory receptor family entry (IPR000725;(Blum et al. 2025). Sequences annotated as pseudogenes or putative pseudogenes were excluded. These reference sequences were aligned with MAFFT v7.505 (Katoh and Standley 2013) using *--localpair* and *--maxiterate 1000*, and the profile was built from this alignment with hmmbuild (HMMER v3.4; (Potter et al. 2018). Predictions that shared ≥ 98% identity were clustered and only the longest retained, and fragments < 30 amino acids were discarded using BITACORA’s internal filters.

ORs belong to the G protein-coupled receptor (GPCR) superfamily, so similarity-based searches can also recover non-OR receptors. We removed these phylogenetically. For each genome, candidate proteins were added with MAFFT (*--add*) to a curated reference alignment of vertebrate ORs and non-OR GPCR outgroups (Policarpo et al. 2024), and a maximum-likelihood tree was inferred with IQ-TREE v.2.2.2.6 (Minh et al. 2020) under the JTT+F+I+G4 model. Each tree was rooted on the outgroup sequences, and candidates falling outside the vertebrate OR clade were discarded.

We next examined the retained candidates for predictions representing fragments of a single locus, merging after phylogenetic filtering so that non-OR sequences could not contribute to merged products. Human ORs are predominantly single-exon genes encoding proteins of approximately 300-360 amino acids, so short predictions occurring close together on the same contig and strand may correspond to one disrupted gene rather than to independent loci. Predictions on the same contig and strand separated by ≤ 150 bp were merged when the combined predicted protein did not exceed 360 amino acids. Merged predictions were directly classified as pseudogenes, as they reconstruct a locus with an internal disruption, and were not subject to the structural classification below. After the merging step, all predicted proteins shorter than 50 amino acids were discarded.

The remaining sequences were classified as intact ORs, pseudogenes or truncated ORs on the basis of their predicted transmembrane domain structure and position within the contig. ORs are seven-transmembrane receptors so the number of predicted transmembrane domains was used as the primary criterion. Domains were predicted with Phobius v.1.01 (Käll et al. 2007) and TMHMM v.2.0c (Krogh et al. 2001), and sequences with at least seven predicted domains under either program were classified as intact. Sequences with fewer than seven predicted domains under both programs were further classified according to their position within the contig. OR coding sequences are approximately 1 kb long, so an incomplete prediction located within 1 kb of a contig boundary may be interrupted by that boundary rather than by a biological gene disruption; these were classified as truncated ORs, whereas incomplete predictions more than 1 kb from both contig ends were classified as pseudogenes. The same 1 kb window defines the truncation event modelled in the following section.

The pseudogene class accordingly includes sequences carrying disrupting mutations, but also sequences retaining a reading frame of approximately full length that are not predicted to form seven transmembrane domains. We treat the latter as non-functional on the grounds that a protein lacking the canonical topology is unlikely to act as an odorant receptor, whether or not its reading frame is interrupted. This is a broader criterion than classifications built on reading-frame integrity and identified loss-of-function mutations (Niimura and Nei 2007; Policarpo et al. 2024). It is applied identically across assemblies, so counts are internally comparable, but they may not be directly comparable with published OR pseudogene counts obtained under the narrower definition.

### Correction of OR counts for assembly fragmentation

Truncated sequences may correspond to an intact OR or to a pseudogene. Discarding them would underestimate repertoire size, but adding them all to the counts inflated it, and did so most in the most fragmented assemblies. The reason is that the number of truncated sequences is likely not the number of loci they represent as, for example, a locus interrupted by a contig boundary can be recovered as two separate predictions. Repertoire size analyses therefore require an estimate of how many loci the truncated sequences stand for, which we obtained with the model described below. Analyses of repertoire composition do not, because each sequence contributes only through the orthogroup it is assigned to, and orthogroups composed entirely of truncated sequences were excluded (see below).

Previous work on primate OR repertoires has corrected repertoire size for fragmentation by regressing the proportion of truncated ORs on inverse N50 (Niimura et al. 2018). In our dataset this relationship was weak (Pearson’s r = 0.22, *p* = 0.06; Spearman’s ⍴ = 0.15, *p* = 0.21; Supplementary Fig. 2). Because N50 is a rough summary of contig length distribution, two assemblies with the same N50 can differ substantially in how much sequence lies near a contig boundary, which is the quantity that determines truncation. Thus, applying the N50-based correction produced implausible repertoire sizes throughout, up to approximatelytwice the count recovered from chromosome-level assemblies in congeners (Supplementary Fig. 3).

We therefore estimated the number of interrupted loci from assembly structure, independently of the observed truncated counts. The model requires an expected number of OR loci per genome, and we take for this the estimated ancestral platyrrhine total repertoire size, rather than each genome’s own observed total, which is itself reduced by fragmentation. This was reconstructed from the eight long-read assemblies with *fastAnc* from phytools R package (Revell 2024) under Brownian motion on log-transformed counts, giving an estimate of 521 loci (N*_OR_;* sum of intact ORs and pseudogenes; 95% CI 471–576). Brownian motion was preferred to an Ornstein–Uhlenbeck model (ΔAICc = 4.78). For each assembly, we calculated the probability that a locus of length *g* = 1 kb, placed at random, overhangs a contig boundary. A gene of length *g* can occupy ℓ + *g* − 1 starting positions relative to a contig of length ℓ, including positions in which the gene extends beyond either contig boundary. Of these, 2(*g* - 1) positions result in truncation at one of the two contig edges. Thus, for an assembly of *n_c_* contigs of total length *G*, the probability of truncation is

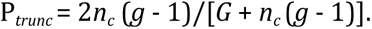

Contigs shorter than 1 kb were excluded from *n_c_* and *G*. The expected number of loci interrupted by assembly boundaries was E(T) = N*_OR_* × P*_trunc_*, and these were added back to the observed counts, divided between intact ORs and pseudogenes in the proportion observed in that genome. Uncertainty in E(T) was obtained by Monte Carlo simulation, sampling 10,000 N_OR_ values from a normal distribution defined by the ancestral reconstruction estimate and its standard deviation, and then drawing truncated counts from a binomial distribution with N_OR_ trials and probability P*_trunc_*. Corrected counts and their intervals are reported in Supplementary Data 1. Functional repertoire size, as used throughout, refers to the corrected number of intact ORs. Because the correction is derived from contig structure and a single ancestral repertoire size applied to all genomes, it uses no information about the taxonomic identity of an assembly. Corrected counts for fragmented assemblies nonetheless converged on those recovered from long-read assemblies of closely related species (Supplementary Fig. 3), so this agreement is independent of the estimation.

### Orthogroup inference and repertoire composition analysis

Orthogroup inference was performed on intact, truncated and pseudogenized OR amino acid sequences with OrthoFinder v.3.1.2 (Emms et al. 2026) with the *-A mafft* option for multiple sequence alignment, and *-T iqtree* for gene-tree inference. We first ran a *core* analysis on the 8 species with high-quality assemblies, and then used the *--assign* option to add the sequences of the remaining species directly to the previous orthogroups or create new orthogroups when appropriate. This allowed us to anchor the composition analyses on these high-quality genomes. Orthogroups were then extracted from the Orthogroups.tsv output file. For subsequent analyses, 31 orthogroups composed exclusively of truncated sequences were excluded.

For the repertoire composition analyses we built a presence-absence matrix based on intact and truncated sequences, using only orthogroups that have at least one intact sequence. An orthogroup is scored as present in a species when at least one of its OR sequences was assigned to it. Retaining truncated sequences can recover orthogroups that would otherwise be scored as absent, but it also risks scoring an orthogroup as present based on a truncated fragment of a pseudogene. If this bias were substantial, fragmented assemblies would be expected to shift away from their long-read relatives in compositional space. We did not observe this pattern: instead, fragmented assemblies clustered with the long-read assembly of their closest relative (Fig. 1b,c). Compositional differences among species were quantified from this matrix as Jaccard dissimilarities computed with the vegan package (Oksanen et al. 2026) for R (R Core Team 2025), and summarized by principal coordinate analyses (PCoA) using the cmdscale base R function. The species phylogeny was projected into the compositional space using the phylomorphospace function from phytools.

### Detection of structural deletions of OR-containing regions in Saimiri boliviensis

To test whether the reduced *Saimiri* repertoire reflects the loss of contiguous OR-containing sequence, we searched for genomic regions containing OR genes in a reference platyrrhine genome but absent from *Saimiri*. We used the chromosome-level assemblies of *C. jacchus* as the reference and *P. cupreus* as an outgroup, allowing regions absent from *Saimiri* to be distinguished from regions gained in *Callithrix*. ORs were annotated in all three genomes with the pipeline described above.

We aligned the *S. boliviensis* assembly to the *C. jacchus* reference with minimap2 v.2.28 (Li 2021) using the asm10 preset, retaining primary chromosomal alignments with mapping quality ≥ 20. Candidate deleted regions were defined as *C. jacchus* intervals containing one or more annotated ORs and overlapping no retained alignment block. Each interval was delimited by the nearest alignment blocks upstream and downstream in reference coordinates, and summarized by position, length, and the number and type of OR genes it contained.

To validate these regions as true absences and rule out assembly or genome-alignment artifacts, we mapped raw PacBio HiFi reads underlying the *S. boliviensis* assembly (SRA accessions SRR32766827-30) to the complete *C. jacchus* reference genome using minimap2 with the *map-hifi* preset. Alignments were filtered to retain reads with mapping quality ≥20, excluding unmapped and secondary alignments. Read depth across each candidate region and its flanking sequence was calculated with bedtools v2.31.1 by averaging coverage across 100 bp windows. Regions with a gap-to-flank coverage ratio below 0.1, calculated using 20 kb flanks on each side, were retained as supported absences.

We then used *P. cupreus* as an outgroup to distinguish losses in *S. boliviensis* from regions specific to *C. jacchus*. The *P. cupreus* assembly was aligned to the *C. jacchus* assembly using the asm10 preset in minimap2 and filtered to retain alignments with mapping quality ≥ 20. Regions depleted in *S. boliviensis* but aligning over at least 80% of their length in *P. cupreus* are present in both a callitrichid and a pitheciid, and were therefore considered as present in the platyrrhine ancestor and lost in *S. boliviensis*. Finally, to distinguish OR loss from relocation elsewhere in the genome, we examined the orthogroup membership of the ORs contained in the validated regions. Each *C. jacchus* OR in these regions was scored as having a potential counterpart in *S. boliviensis* when at least one *S. boliviensis* sequence was assigned to the same orthogroup.

### OR ancestral retention and focal duplication

To quantify ancestral OR retention, we inferred which OR orthogroups were present in the last common ancestor of platyrrhines based on the presence of intact OR copies in the 80 sampled species, assuming Dollo parsimony. We then assessed the fate of these ancestral orthogroups in each of the eight species with high-quality genome assemblies. Each orthogroup was classified as retained if at least one intact OR copy was present, pseudogenized if only pseudogene copies remained, or absent if no sequence belonging to that orthogroup was detected (truncated sequences are nearly absent in these high-quality assemblies).

Finally, we searched for recent focal OR expansions in capuchins and *Aotus* using a phylogeny of intact OR proteins from the long-read genomes of *A. nancymaae*, *C. albifrons*, *S. apella*, together with *C. jacchus and S. boliviensis* as representatives of their respective sister lineages, plus *B. arachnoides* as an outgroup. Sequences were aligned with MAFFT v.7.505 using the FFT-NS-2 strategy, and trimmed with trimAl v.1.5 (Capella-Gutiérrez et al. 2009) with a minimum site occupancy of 50% (*-gt* 0.5). A less stringent threshold (*-gt* 0.2) produced almost identical results (not shown). Maximum-likelihood phylogenies were inferred with IQ-TREE v.2.2.2.6 using ModelFinder and 1,000 ultrafast bootstrap replicates. Clades supported by bootstrap ≥ 95% and composed exclusively of two or more sequences from *A. nancymaae*, or three or more from the two capuchins, were taken as evidence of duplication; two capuchin sequences are expected under simple orthology.

### Selection analyses on OR coding sequences

To characterize selective pressures acting on OR coding sequences, we analysed one species per genus, using in each case the genome assembly with the highest N50. Sampling one representative per genus rather than all 80 species means that each terminal branch spans the interval since divergence from its sister genus, so the branches tested are expected to be long enough to accumulate detectable signal and the results describe selection over the history of each genus rather than only its most recent portion.

Codon alignments were built from intact and truncated sequences of each orthogroup, excluding orthogroups with fewer than five sequences. Coding sequences were extracted from the assemblies using the OR annotations, and aligned with PAL2NAL (Suyama et al. 2006) using the corresponding OrthoFinder protein alignments as templates. Orthogroups with more than one sequence that failed to align were removed; those with only one sequence that failed to align were maintained but the problematic sequence was removed. Gene trees were taken from the resolved gene trees produced by OrthoFinder and pruned to match the sequences in each alignment. After filtering, 603 orthogroups were retained for analysis.

Branch-specific dN/dS ratios (ω) were estimated with the free-ratio model in CODEML (PAML v. 4.10.10; (Yang 2007), fitted independently to each orthogroup with model = 1, NSsites = 0, and clock = 0, and with gene trees treated as unrooted. Summaries were restricted to terminal branches. Estimates with dS = 0 or with ω at the upper optimization bound (ω = 999) were classified as non-valid and excluded from downstream analyses. All remaining estimates were considered valid. Estimates at the lower bound (ω = 0.0001) were retained when classifying branches relative to ω = 1, but their magnitude was not interpreted and they are omitted from Fig. 4a.

Episodic positive selection was tested with the adaptive branch-site random effects likelihood model (aBSREL v2.6; (M. D. Smith et al. 2015) implemented in HyPhy v2.5.96, run independently for each orthogroup across all branches of the corresponding gene tree, with subsequent analyses focusing only on terminal branches. Because significance had to be assessed across orthogroups as well as across branches, uncorrected P values from all tested terminal branches were pooled across orthogroups and adjusted together with the Benjamini–Hochberg procedure, at a false discovery rate of 0.05. As a sensitivity analysis, we restricted the aBSREL results to terminal branches corresponding to intact ORs only. This retained 598 orthogroups and recovered the same overall pattern (Supplementary Fig. 6).

### Morphological sampling and trait quantification

We quantified olfaction-related cranial morphology from computed tomography (CT) and µCT scans of 59 adult specimens representing 33 platyrrhine species, 19 genera and all five families. Both sexes were included whenever possible. Because our aim was to examine macroevolutionary variation across species rather than intraspecific variation, sampling prioritized taxonomic breadth. µCT scans were obtained from an existing dataset previously assembled by our research group from different museums (Aristide et al. 2015) and CT scans from the online repositories MorphoSource and the Digital Morphology Museum of the Kyoto University Primate Research Institute (KUPRI). Differences in resolution among CT sources are not expected to affect measurements as they are smaller than interspecific variation (Aristide et al. 2015). The source and information for each specimen are provided in Supplementary Data 5.

We measured four olfaction-related traits: olfactory bulb endocast volume (OBV), cribriform plate area (CPA), olfactory turbinal surface area (TSA) and turbinal number. Segmentations and measurements were performed by J. B. Romero in 3D Slicer v.5.10 (Fedorov et al. 2012) using the Segment Editor, Segment Statistics and SlicerMorph (Rolfe et al. 2021) modules. OBV was estimated by generating a cranial endocast and manually isolating its olfactory bulb portion, including the accessory olfactory bulb, which is extremely small in platyrrhines (Laska and Salazar 2015). The posterior limit of the bulb was defined in the sagittal plane at the point where the cranial endocast curvature changed from concave to convex. Thus, OBV refers to the volume of the olfactory bulb endocast rather than neural tissue itself. Endocranial volume (ECV) was calculated using the full cranial endocast. The olfactory turbinals (nasoturbinal and ethmoturbinals I-III) were manually segmented on the single, best-preserved side of each specimen, following Lundeen and Kay (2022). CPA was calculated by manually tracing the cribriform plate contour using the Closed Curve tool in 3D Slicer, whereas TSA was calculated as the summed surface area of the olfactory turbinals. Turbinal number was defined as the number of distinct ethmoturbinals present on the segmented side, as the nasoturbinal was always present. 27 species were represented by two specimens, and individual measurements were averaged to obtain a species-level value, while six species were represented by a single individual. Individual measurements and the species-level dataset used in subsequent analyses are provided in Supplementary Data 5.

### Ecological data

We compiled species-level information on diet, activity pattern, color vision and body mass from published sources (Supplementary Data 2 and references therein; Supplementary Fig. 4). Diet was quantified as the percentage of feeding time devoted to each of five food categories, fruits, leaves, seeds, animal prey, and exudates, rather than by assigning species to broad dietary categories, thereby retaining variation among closely related species. Each dietary component was analysed separately as these proportions are not statistically independent. Dietary data were available for 51 species, except for exudate consumption, which was available for 50. Activity pattern was classified as diurnal or non-diurnal, the latter including nocturnal and cathemeral species. Color vision was scored following (Niimura et al. 2018), on an ordinal scale ranking color vision categories by discriminative capacity: monochromatic (0), polymorphic dichromatic–trichromatic (2) and routinely trichromatic (3) species. No platyrrhine is routinely dichromatic, so score 1 is not represented in our sample. We retained the scheme rather than collapsing the scale, so that the interval separating monochromatic species from those with polymorphic trichromatism is larger than that separating polymorphic from the routine trichromatic species.

### Phylogenetic comparative analyses

We used phylogenetic generalized least-squares models (PGLS) to test whether OR repertoire size covaried with ecological and sensory variables, or olfaction-related morphological traits, while accounting for shared ancestry. Models were fitted in R using the phylolm package (Ho and Ané 2014), with Pagel’s λ estimated by maximum likelihood. OR counts and continuous morphological variables were natural log-transformed before model fitting.

For ecological analyses (n = 28-51 species), corrected functional and pseudogene OR counts were tested separately against each dietary variable, color vision (included as an ordinal predictor) or activity pattern. We also fitted models including frugivory or folivory together with activity pattern or color vision, to assess whether a dietary association emerged once this variation was accounted for.

For morphological analyses (n = 33 species), we first regressed functional OR number on OBV and CPA with body mass (BM) and ECV as covariates, as platyrrhines vary both in body size and encephalization. For regressions against TSA, only body mass was included as covariate, as ECV is not expected to be directly related to TSA, while for turbinal number, no allometric scaling variable was included. We then focused on OBV, the trait most strongly associated with repertoire size, and fitted three models: OBV on BM and ECV (relative bulb size); OBV on functional OR number, BM, and ECV (bulb size relative to repertoire size); and functional OR number on OBV, BM and ECV (repertoire size relative to bulb size). We fitted the relationship in both directions because, with an imperfect correlation, a lineage that departs from the shared trend in one trait deviates most clearly when that trait is the response; the two directions were therefore interpreted jointly.

To test whether particular lineages departed from these relationships, we added binary terms separately for *Saimiri* and *Aotus* (1 for species of the genus, 0 otherwise). For the genomic model relating pseudogene number to functional OR number (n = 80), the model including the *Saimiri* term was compared with the model without it using a likelihood-ratio test. For the morphological models, genus effects were assessed from the coefficient of the binary term. To visualize where on the phylogeny departures arose, residuals from the three OBV models, fitted without genus terms, were standardized (z-scores) and their evolution reconstructed under Brownian motion with the contMap function in phytools. These reconstructions are presented for visualization only; departures are quantified by the binary terms.

Benjamini–Hochberg correction was applied separately to two families of univariate ecological tests: diet and activity–color vision. P-values from morphological and multivariate models are reported without multiple-testing correction. Complete model results are reported in Supplementary Tables 2, 3 and 8.

