## Supplementary Material for "Lineage-specific histories rather than shared drivers shape olfactory evolution in New World monkeys"

**[2]CONICET. Museo Histórico y Arqueológico "Ricardo Pascual Rosa", Neuquén, Argentina.**

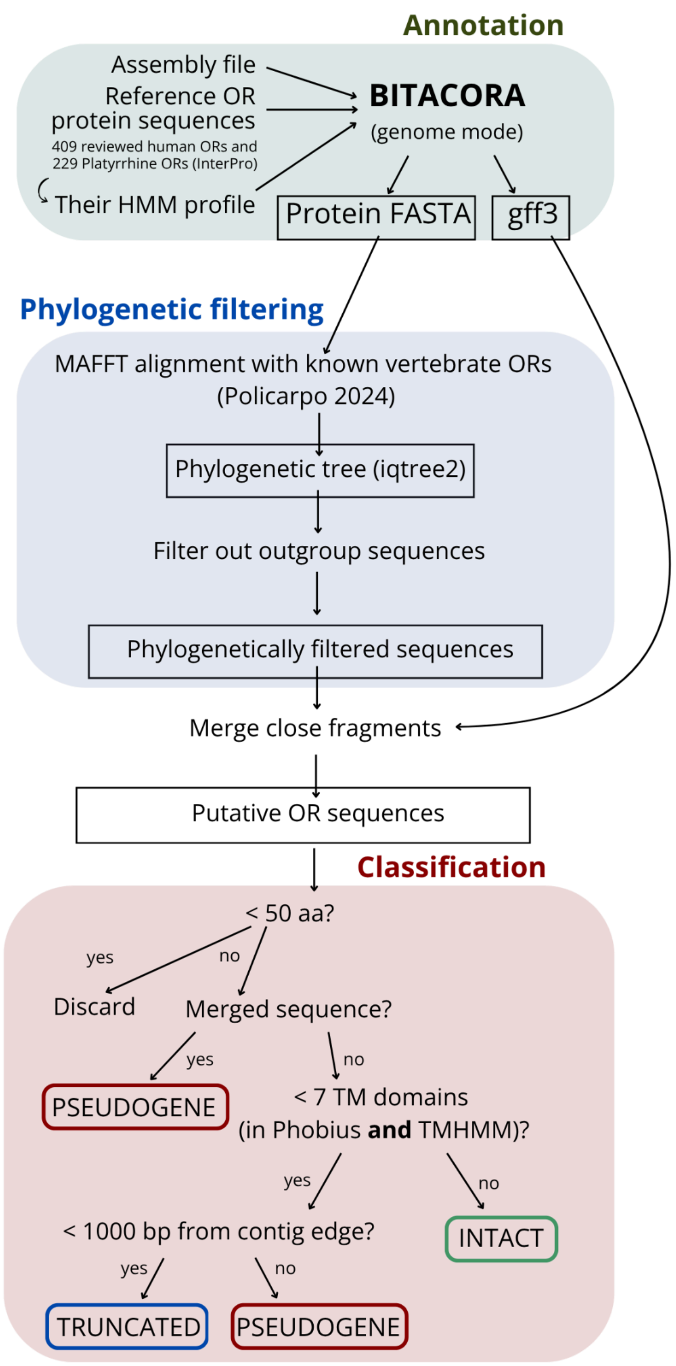


**Supplementary Figure 1. Scheme of the olfactory receptor (OR) gene annotation and classification pipeline.** Candidate OR sequences identified with BITACORA were phylogenetically filtered, and adjacent fragments were merged when appropriate. Sequences shorter than 50 amino-acids were discarded. Retained sequences were classified as intact, pseudogenized or truncated based on fragment merging, predicted transmembrane (TM) domains and proximity to contig boundaries (see Methods).

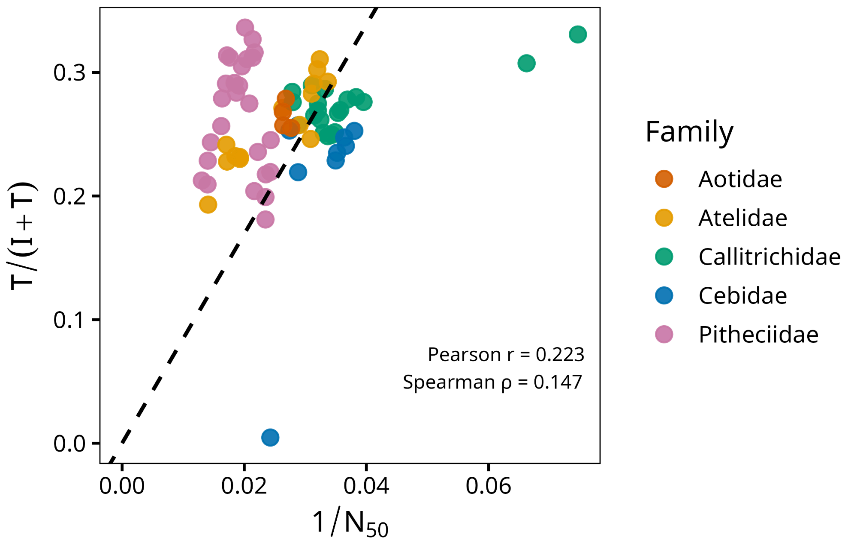


**Supplementary Figure 2 - Weak association between Contig N50 and the proportion of truncated olfactory receptor sequences in the 72 fragmented assemblies.** Each point represents a species, colored by family. The proportion of truncated OR sequences, calculated as T/(I + T), where I and T denote observed intact and truncated sequences, respectively, is plotted against inverse contig N50 (N50 expressed in kb). Pearson's and Spearman's correlation coefficients are shown. The dashed line represents a linear regression constrained to pass through the origin and fitted to the same 72 species.

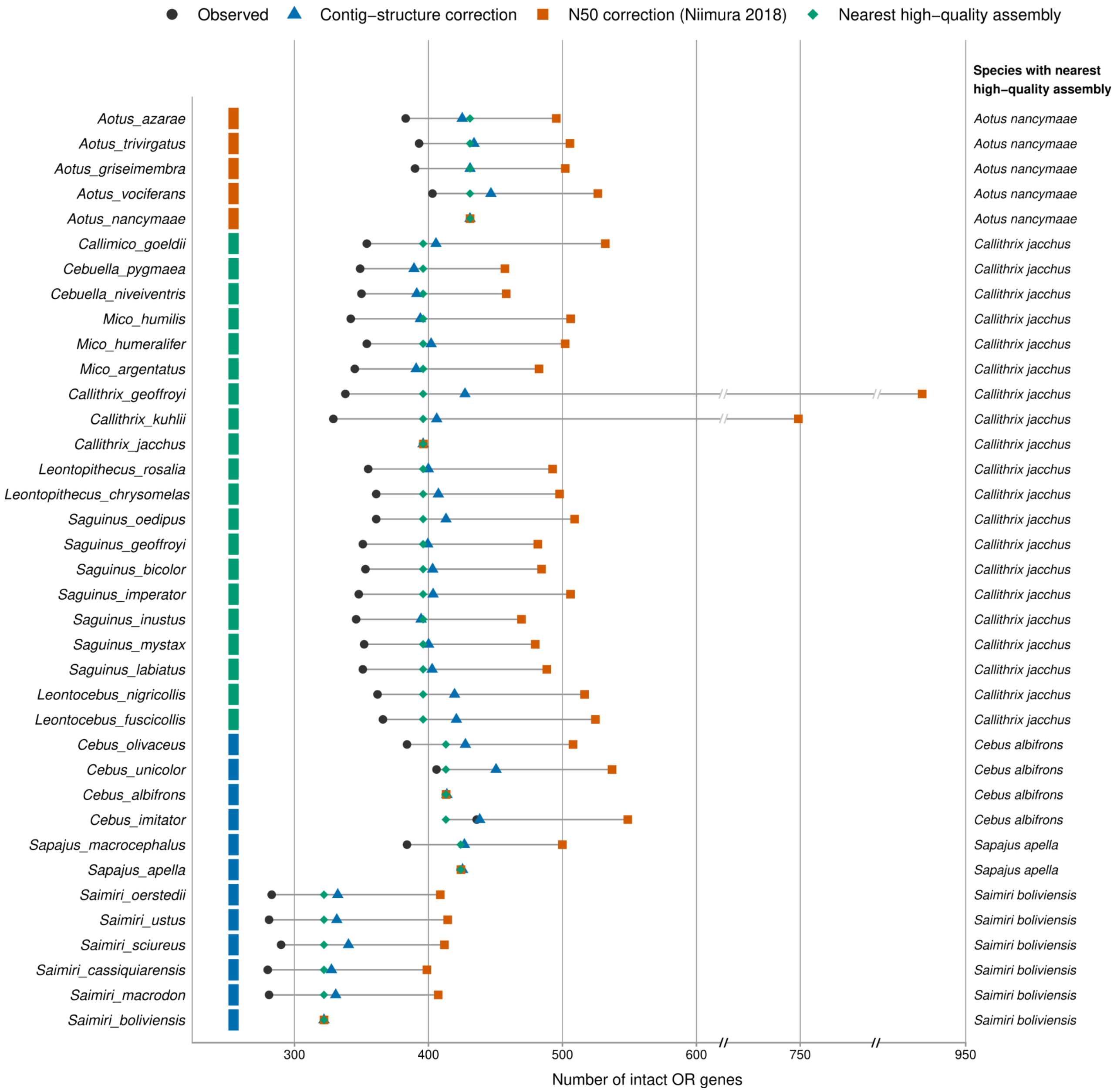


**Supplementary Figure 3. Comparison of intact OR counts under alternative corrections for assembly fragmentation.** Symbols indicate observed counts (black dots), counts corrected using contig structure as used in this study (blue triangles), counts corrected using the N50-based approach of Niimura et al. (2018; orange squares), and counts in the nearest phylogenetic relative with a high quality assembly (green diamonds). For each species, the nearest phylogenetic relative with a high quality assembly is listed on the right. Colour strips indicate families, as in Fig. 1. Note the breaks in the horizontal axis.

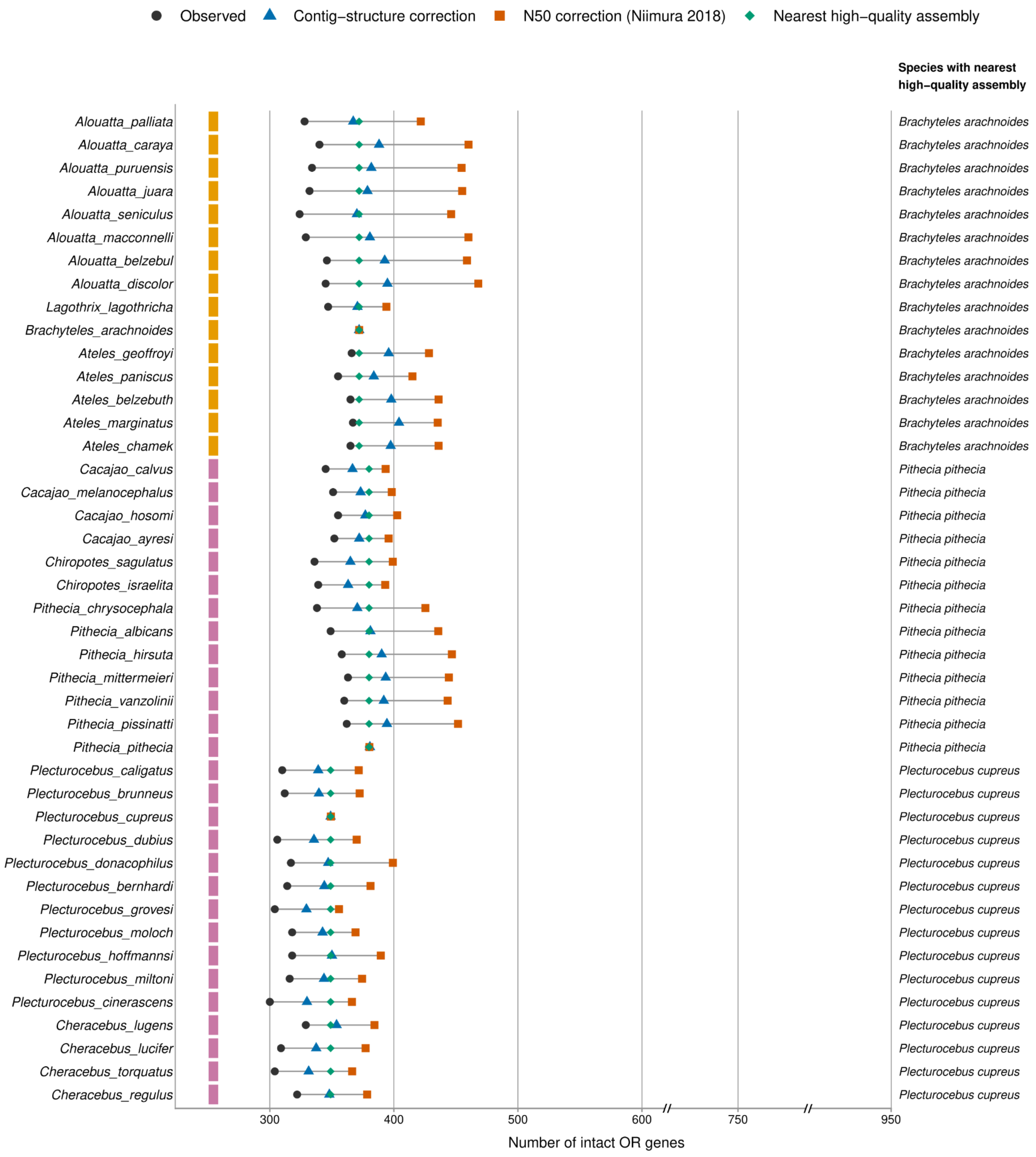


**Supplementary Figure 3 (continued).**

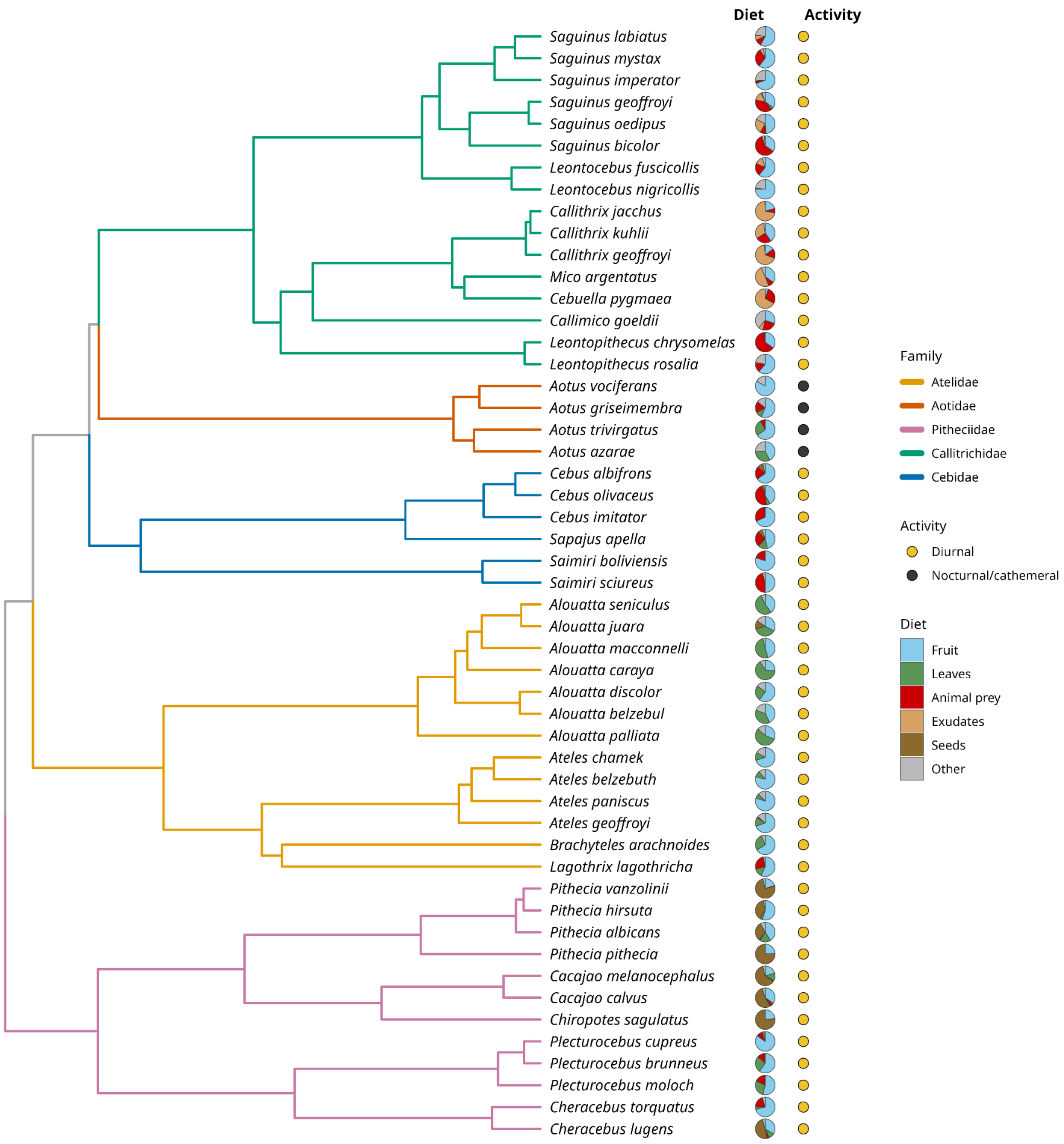


**Supplementary Figure 4 - Dietary composition and activity patterns across platyrrhines.** Pie charts show the proportion of feeding time devoted to each food category. Circles indicate diurnal (yellow) or nocturnal/cathemeral (dark grey) activity. Phylogenetic branches are coloured by family. Only species with available dietary data are shown.

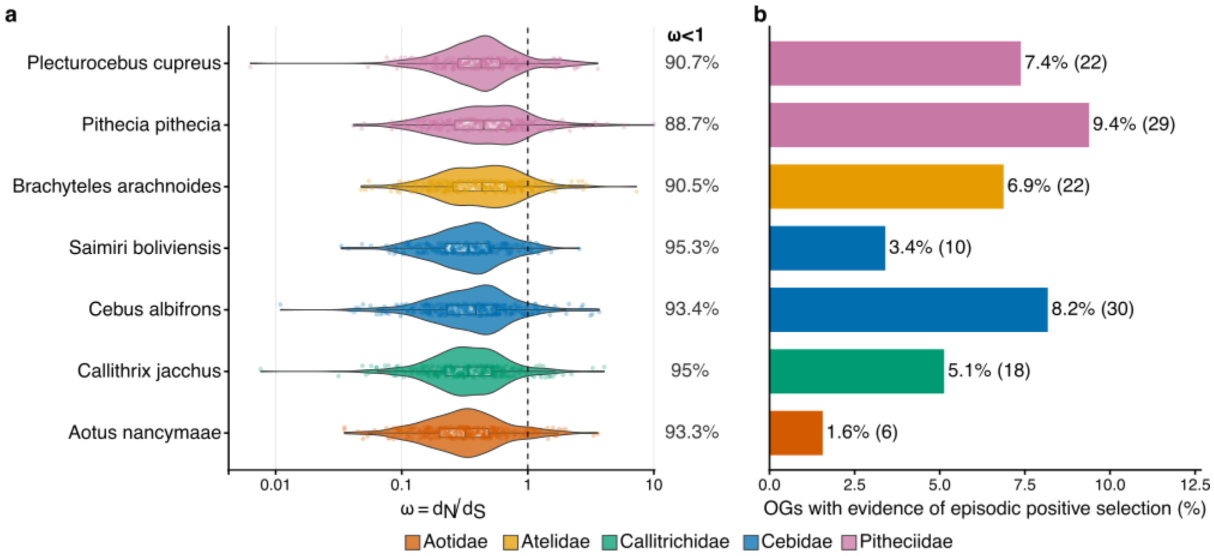


**Supplementary Figure 5. Selection analyses of OR genes in seven species with high-quality genome assemblies.** Only one capuchin species (*Cebus albifrons*) was included to keep terminal branch lengths comparable. **a.** Distributions of terminal branch-specific ω (dN/dS) estimated using CODEML, shown on a logarithmic scale. The dashed line marks ω = 1; percentages indicate the proportion of branches with ω < 1. Colours indicate families. **b.** Percentage of tested orthogroups with evidence of episodic positive selection in each species (aBSREL, BH-adjusted $p < 0.05$); numbers in parentheses indicate significant orthogroup counts.

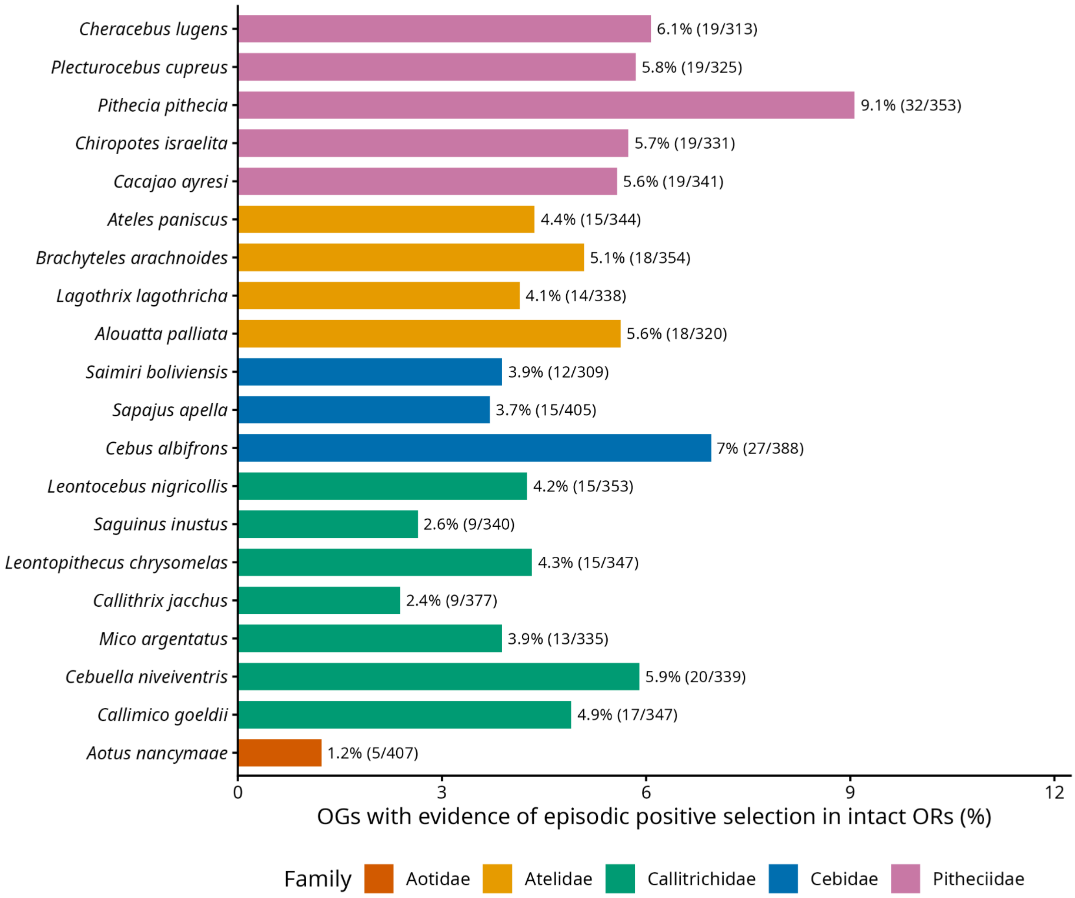


**Supplementary Figure 6. aBSREL selection analysis results excluding truncated ORs.** Percentage of tested orthogroups with evidence of episodic positive selection in each species (aBSREL, BH-adjusted $p < 0.05$); numbers in parentheses indicate significant orthogroup counts.

### Supplementary tables

**Supplementary Table 1. Genome assemblies included in this study.**
Assemblies in bold are long-read based (high-quality). NCBI IDs are provided.

| ID | Species | Family | Contig N50 (bp) |
|---|---|---|---|
| GCA_963573465.1 | *Aotus azarae* | Aotidae | 37300 |
| GCA_963574505.1 | *Aotus griseimembra* | Aotidae | 37900 |
| **GCF_030222135.1** | ***Aotus nancymaae*** | **Aotidae** | **55000000** |
| GCA_963574785.1 | *Aotus trivirgatus* | Aotidae | 38000 |
| GCA_963573775.1 | *Aotus vociferans* | Aotidae | 36100 |
| GCA_963574145.1 | *Alouatta belzebul* | Atelidae | 34400 |
| GCA_963575295.1 | *Alouatta caraya* | Atelidae | 32400 |
| GCA_963575205.1 | *Alouatta discolor* | Atelidae | 32200 |
| GCA_963574365.1 | *Alouatta juara* | Atelidae | 31300 |
| GCA_963573945.1 | *Alouatta macconnelli* | Atelidae | 29700 |
| GCA_963573975.1 | *Alouatta palliata* | Atelidae | 38100 |
| GCA_963574235.1 | *Alouatta puruensis* | Atelidae | 31900 |
| GCA_963573765.1 | *Alouatta seniculus* | Atelidae | 30900 |
| GCA_963573605.1 | *Ateles belzebuth* | Atelidae | 51900 |
| GCA_963574935.1 | *Ateles chamek* | Atelidae | 51900 |
| GCA_963574205.1 | *Ateles geoffroyi* | Atelidae | 58200 |
| GCA_963573555.1 | *Ateles marginatus* | Atelidae | 53900 |
| GCA_963573985.1 | *Ateles paniscus* | Atelidae | 58600 |
| **GCA_047496155.1** | ***Brachyteles arachnoides*** | **Atelidae** | **58900000** |
| GCA_963574225.1 | *Lagothrix lagothricha* | Atelidae | 71000 |
| GCA_963573925.1 | *Callimico goeldii* | Callitrichidae | 25300 |
| GCA_963573515.1 | *Callithrix geoffroyi* | Callitrichidae | 13400 |
| **GCF_011100555.1** | ***Callithrix jacchus*** | **Callitrichidae** | **13200000** |
| GCA_963573495.1 | *Callithrix kuhlii* | Callitrichidae | 15100 |
| GCA_963575025.1 | *Cebuella niveiventris* | Callitrichidae | 35900 |
| GCA_963575105.1 | *Cebuella pygmaea* | Callitrichidae | 35800 |
| GCA_963575285.1 | *Leontocebus fuscicollis* | Callitrichidae | 28000 |
| GCA_963575265.1 | *Leontocebus nigricollis* | Callitrichidae | 28300 |
| GCA_963574395.1 | *Leontopithecus chrysomelas* | Callitrichidae | 30800 |
| GCA_963574445.1 | *Leontopithecus rosalia* | Callitrichidae | 30300 |
| GCA_963573565.1 | *Mico argentatus* | Callitrichidae | 29700 |
| GCA_963574605.1 | *Mico humeralifer* | Callitrichidae | 28700 |
| GCA_963574545.1 | *Mico humilis* | Callitrichidae | 26100 |
| GCA_963574455.1 | *Saguinus bicolor* | Callitrichidae | 31200 |
| GCA_963574215.1 | *Saguinus geoffroyi* | Callitrichidae | 31200 |
| GCA_963574955.1 | *Saguinus imperator* | Callitrichidae | 27100 |
| GCA_963574735.1 | *Saguinus inustus* | Callitrichidae | 32200 |
| GCA_963575005.1 | *Saguinus labiatus* | Callitrichidae | 30100 |
| GCA_963574345.1 | *Saguinus mystax* | Callitrichidae | 31800 |

| ID | Species | Family | Contig N50 (bp) |
|---|---|---|---|
| GCA_963574915.1 | *Saguinus oedipus* | Callitrichidae | 29100 |
| **GCA_023783575.1** | ***Cebus albifrons*** | **Cebidae** | **18500000** |
| GCF_001604975.1 | *Cebus imitator* | Cebidae | 41200 |
| GCA_963574745.1 | *Cebus olivaceus* | Cebidae | 34700 |
| GCA_963574255.1 | *Cebus unicolor* | Cebidae | 34700 |
| **GCF_048565385.1** | ***Saimiri boliviensis*** | **Cebidae** | **69222429** |
| GCA_963573395.1 | *Saimiri cassiquiarensis* | Cebidae | 28400 |
| GCA_963573505.1 | *Saimiri macrodon* | Cebidae | 27300 |
| GCA_963574195.1 | *Saimiri oerstedii* | Cebidae | 27500 |
| GCA_963573345.1 | *Saimiri sciureus* | Cebidae | 28600 |
| GCA_963574015.1 | *Saimiri ustus* | Cebidae | 26300 |
| **GCA_023762875.2** | ***Sapajus apella*** | **Cebidae** | **12600000** |
| GCA_963574755.1 | *Sapajus macrocephalus* | Cebidae | 36500 |
| GCA_963573425.1 | *Cacajao ayresi* | Pitheciidae | 76600 |
| GCA_963573955.1 | *Cacajao calvus* | Pitheciidae | 68800 |
| GCA_963573795.1 | *Cacajao hosomi* | Pitheciidae | 71500 |
| GCA_963573385.1 | *Cacajao melanocephalus* | Pitheciidae | 71300 |
| GCA_963573755.1 | *Cheracebus lucifer* | Pitheciidae | 46800 |
| GCA_963574535.1 | *Cheracebus lugens* | Pitheciidae | 58800 |
| GCA_963574635.1 | *Cheracebus regulus* | Pitheciidae | 56700 |
| GCA_963574645.1 | *Cheracebus torquatus* | Pitheciidae | 49700 |
| GCA_963573965.1 | *Chiropotes israelita* | Pitheciidae | 61600 |
| GCA_963574175.1 | *Chiropotes sagulatus* | Pitheciidae | 53500 |
| GCA_963573695.1 | *Pithecia albicans* | Pitheciidae | 42500 |
| GCA_963574905.1 | *Pithecia chrysocephala* | Pitheciidae | 41200 |
| GCA_963573645.1 | *Pithecia hirsuta* | Pitheciidae | 42600 |
| GCA_963574975.1 | *Pithecia mittermeieri* | Pitheciidae | 46200 |
| GCA_963575305.1 | *Pithecia pissinatti* | Pitheciidae | 42600 |
| **GCA_023779675.1** | ***Pithecia pithecia*** | **Pitheciidae** | **10900000** |
| GCA_963574005.1 | *Pithecia vanzolinii* | Pitheciidae | 45000 |
| GCA_963574705.1 | *Plecturocebus bernhardi* | Pitheciidae | 48000 |
| GCA_963573885.1 | *Plecturocebus brunneus* | Pitheciidae | 52200 |
| GCA_963575255.1 | *Plecturocebus caligatus* | Pitheciidae | 51000 |
| GCA_963573415.1 | *Plecturocebus cinerascens* | Pitheciidae | 46800 |
| **GCA_040437455.1** | ***Plecturocebus cupreus*** | **Pitheciidae** | **17000000** |
| GCA_004027715.1 | *Plecturocebus donacophilus* | Pitheciidae | 41100 |
| GCA_963575235.1 | *Plecturocebus dubius* | Pitheciidae | 48900 |
| GCA_963574625.1 | *Plecturocebus grovesi* | Pitheciidae | 58200 |
| GCA_963575245.1 | *Plecturocebus hoffmannsi* | Pitheciidae | 46100 |
| GCA_963575275.1 | *Plecturocebus miltoni* | Pitheciidae | 54300 |
| GCA_963574105.1 | *Plecturocebus moloch* | Pitheciidae | 61100 |

**Supplementary Table 2. Results of PGLS models between functional and pseudogenized OR repertoire size and ecological variables.** Dietary variables are expressed as the percentage of feeding time dedicated to each food category. Activity pattern was coded as diurnal vs non-diurnal, and color vision was analyzed as an ordinal phenotype (0 for monochromatic, 2 for polymorphic and 3 for trichromatic). n indicates the number of species included in each model; β is the regression slope, p indicates the p-value associated with each predictor coefficient, $R^2$ is the model R-squared, and lambda is the Pagel's phylogenetic signal estimated for each model. For univariate models, p-values are Benjamini–Hochberg adjusted. For multivariate models, nominal p-values are reported. Bold p-values indicate p < 0.05. For the high-quality genome sensitivity analysis, missing dietary values for *Aotus nancymaae* were replaced by the corresponding mean across other Aotus species. Missing color-vision scores were assigned according to congeners (A. nancymaae: 0; Cebus albifrons: 2).

| Dataset | Univariate model | n | Predictor | slope (β) | BH adjusted p | R2 | λ |
|---|---|---|---|---|---|---|---|
| All species with available data | log(ORs) ~ %Fruit | 51 | %Fruit | 0.000 | 0.886 | 0.005 | 0.951 |
| | log(ORs) ~ %Leaves | 51 | %Leaves | -0.001 | 0.750 | 0.035 | 0.956 |
| | log(ORs) ~ %Animal_prey | 51 | %Animal prey | 0.000 | 0.886 | 0.000 | 0.949 |
| | log(ORs) ~ %Seeds | 51 | %Seeds | 0.000 | 0.750 | 0.030 | 0.955 |
| | log(ORs) ~ %Exudates | 50 | %Exudates | 0.000 | 0.886 | 0.002 | 0.949 |
| | log(psg) ~ %Fruit | 51 | %Fruit | -0.001 | 0.886 | 0.017 | 0.855 |
| | log(psg) ~ %Leaves | 51 | %Leaves | 0.002 | 0.750 | 0.039 | 0.896 |
| | log(psg) ~ %Animal_prey | 51 | %Animal prey | 0.000 | 0.886 | 0.001 | 0.873 |
| | log(psg) ~ %Seeds | 51 | %Seeds | -0.001 | 0.886 | 0.005 | 0.889 |
| | log(psg) ~ %Exudates | 50 | %Exudates | -0.001 | 0.886 | 0.003 | 0.879 |
| | log(ORs) ~ Color vision | 30 | Color vision | -0.048 | 0.078 | 0.107 | 0.911 |
| | log(psg) ~ Color vision | 30 | Color vision | 0.157 | **0.018** | 0.221 | 0.69 |
| | log(ORs) ~ Activity | 80 | Activity | -0.121 | **0.041** | 0.058 | 0.932 |
| | log(psg) ~ Activity | 80 | Activity | 0.347 | **0.008** | 0.115 | 0.754 |
| Species with high-quality genomes | log(ORs) ~ %Fruit | 8 | %Fruit | -0.002 | 0.825 | 0.248 | 1 |
| | log(ORs) ~ %Leaves | 8 | %Leaves | 0.003 | 0.825 | 0.142 | 1 |
| | log(ORs) ~ %Animal_prey | 8 | %Animal prey | 0.001 | 0.825 | 0.009 | <0.001 |
| | log(ORs) ~ %Seeds | 8 | %Seeds | 0.000 | 0.825 | 0.011 | 1 |
| | log(ORs) ~ %Exudates | 8 | %Exudates | 0.000 | 0.825 | 0.014 | <0.001 |
| | log(psg) ~ %Fruit | 8 | %Fruit | -0.005 | 0.825 | 0.209 | 1 |
| | log(psg) ~ %Leaves | 8 | %Leaves | -0.004 | 0.825 | 0.041 | 1 |
| | log(psg) ~ %Animal_prey | 8 | %Animal prey | -0.006 | 0.825 | 0.034 | 1 |
| | log(psg) ~ %Seeds | 8 | %Seeds | 0.006 | 0.825 | 0.351 | <0.001 |
| | log(psg) ~ %Exudates | 8 | %Exudates | 0.002 | 0.825 | 0.027 | 1 |
| | log(ORs) ~ Color vision | 7 | Color vision | -0.058 | 0.324 | 0.193 | <0.001 |
| | log(psg) ~ Color vision | 7 | Color vision | 0.190 | 0.310 | 0.270 | 1 |
| | log(ORs) ~ Activity | 8 | Activity | -0.137 | 0.310 | 0.230 | 1 |
| | log(psg) ~ Activity | 8 | Activity | 0.386 | 0.310 | 0.277 | 1 |

| Dataset | Multivariate model | n | Predictor | slope (β) | p | R2 | λ |
|---|---|---|---|---|---|---|---|
| All species with available data | log(ORs) ~ %Fruit + Activity | 51 | %Fruit | 0.000 | 0.553 | 0.086 | 0.938 |
| | | | Activity | -0.125 | **0.044** | | |
| | log(ORs) ~ %Leaves + Activity | 51 | %Leaves | -0.001 | 0.149 | 0.117 | 0.943 |
| | | | Activity | -0.129 | **0.036** | | |
| | log(ORs) ~ %Animal_prey + Activity | 51 | %Animal prey | 0.000 | 0.965 | 0.080 | 0.936 |
| | | | Activity | -0.122 | **0.047** | | |
| | log(ORs) ~ %Seeds + Activity | 51 | % Seeds | 0.001 | 0.192 | 0.110 | 0.944 |
| | | | Activity | -0.127 | **0.040** | | |
| | log(ORs) ~ %Exudates + Activity | 50 | %Exudates | 0.000 | 0.671 | 0.084 | 0.936 |
| | | | Activity | -0.124 | **0.045** | | |
| | log(ORs) ~ %Fruit + Color vision | 28 | %Fruit | -0.001 | 0.206 | 0.153 | 0.908 |
| | | | Color vision | -0.053 | 0.075 | | |
| | log(ORs) ~ %Leaves + Color vision | 28 | %Leaves | 0.000 | 0.643 | 0.105 | 0.902 |
| | | | Color vision | -0.048 | 0.105 | | |
| | log(ORs) ~ %Animal_prey + Color vision | 28 | %Animal prey | 0.000 | 0.838 | 0.098 | 0.905 |
| | | | Color vision | -0.047 | 0.113 | | |
| | log(ORs) ~ %Seeds + Color vision | 28 | % Seeds | 0.000 | 0.716 | 0.102 | 0.901 |
| | | | Color vision | -0.047 | 0.109 | | |
| | log(ORs) ~ %Exudates + Color vision | 28 | %Exudates | 0.000 | 0.668 | 0.105 | 0.898 |
| | | | Color vision | -0.047 | 0.106 | | |
| | log(psg) ~ %Fruit + Activity | 51 | %Fruit | -0.001 | 0.335 | 0.127 | 0.800 |
| | | | Activity | 0.281 | **0.022** | | |
| | log(psg) ~ %Leaves + Activity | 51 | %Leaves | 0.002 | 0.135 | 0.139 | 0.844 |
| | | | Activity | 0.307 | **0.018** | | |
| | log(psg) ~ %Animal_prey + Activity | 51 | %Animal prey | 0.000 | 0.916 | 0.103 | 0.823 |
| | | | Activity | 0.288 | **0.024** | | |
| | log(psg) ~ %Seeds + Activity | 51 | % Seeds | 0.000 | 0.696 | 0.102 | 0.838 |
| | | | Activity | 0.293 | **0.025** | | |
| | log(psg) ~ %Exudates + Activity | 50 | %Exudates | -0.001 | 0.640 | 0.108 | 0.826 |
| | | | Activity | 0.295 | **0.022** | | |
| | log(psg) ~ %Fruit + Color vision | 28 | %Fruit | -0.002 | 0.174 | 0.218 | 0.796 |
| | | | Color vision | 0.12 | 0.061 | | |
| | log(psg) ~ %Leaves + Color vision | 28 | %Leaves | 0.001 | 0.79 | 0.154 | 0.819 |
| | | | Color vision | 0.133 | **0.048** | | |
| | log(psg) ~ %Animal_prey + Color vision | 28 | %Animal prey | 0.000 | 0.759 | 0.154 | 0.824 |
| | | | Color vision | 0.136 | **0.044** | | |
| | log(psg) ~ %Seeds + Color vision | 28 | % Seeds | 0.003 | 0.170 | 0.223 | 0.780 |
| | | | Color vision | 0.131 | **0.036** | | |
| | log(psg) ~ %Exudates + Color vision | 28 | %Exudates | 0.000 | 0.984 | 0.152 | 0.819 |
| | | | Color vision | 0.135 | **0.045** | | |

**Supplementary Table 3. Phylogenetic generalized least squares regressions between functional and pseudogenized OR counts.** PGLS models were fitted with and without a binary *Saimiri* term to test whether *Saimiri* departs from the general cross-species relationship. Negative *Saimiri* coefficients indicate lower values than expected after accounting for the predictors included in each model. λ denotes the estimated phylogenetic signal. ΔAIC is reported relative to the best-fitting model within each pair, and LRT p-values compare models with and without the *Saimiri* term. Analyses restricted to species with high-quality genomes (Contig N50 > 10 Mb) are shown as sensitivity analyses.

| Dataset | Model | n | $R^2$ | λ | Predictor | β | p | Δ AIC | LRT p |
|---|---|---|---|---|---|---|---|---|---|
| All species | log(psg) ~ log(ORs) | 80 | 0.051 | 0.909 | log(ORs) | -0.653 | **0.043** | 2.868 | |
| | log(psg) ~ log(ORs) + *Saimiri* | 80 | 0.107 | 0.852 | log(ORs) | -0.954 | **0.006** | 0 | **0.027** |
| | | | | | *Saimiri* | -0.342 | **0.024** | | |
| Species with high-quality genomes * | log(psg) ~ log(ORs) | 8 | 0.092 | 1 | log(ORs) | 0.778 | 0.465 | 3.334 | |
| | log(psg) ~ log(ORs) + *Saimiri* | 8 | 0.534 | 1 | log(ORs) | -1.385 | 0.324 | 0 | **0.021** |
| | | | | | *Saimiri* | -0.766 | 0.081 | | |

**Supplementary Table 4. Candidate OR-containing deletions in the *Saimiri boliviensis* genome.** Gap coordinates are given relative to the *Callithrix jacchus* reference assembly. OR counts in each gap are reported as intact/pseudogenized ORs (i/p). The gap/flank ratio is the ratio between mean *S. boliviensis* raw-read coverage within each candidate region and mean coverage in its 20 kb flanking regions, obtained by mapping *S. boliviensis* reads to the *C. jacchus* reference genome. *P. cupreus* aligned (%) indicates the percentage of the corresponding *C. jacchus* interval aligned by the *Plecturocebus cupreus* genome. Candidate regions were classified as "Unsupported deletion" when the gap/flank ratio was ≥0.1, "Putative gain in *C. jacchus*" when the ratio was <0.1 but <80% of the interval aligned to *P. cupreus*, and "Supported deletion" when the ratio was <0.1 and ≥80% of the interval aligned to *P. cupreus*. Alignment % was not evaluated for *P. cupreus* when the gap/flank ratio was > 0.1.

| Candidate | *C. jacchus* chr. | Start position | End position | Length (kb) | ORs (i/p) | Gap/flank ratio | *P. cupreus* aligned (%) | Outcome |
|---|---|---|---|---|---|---|---|---|
| gap_01 | NC_071442.1 | 69214291 | 69268242 | 54 | 1/2 | 0.139 | - | Unsupported deletion |
| gap_02 | NC_071442.1 | 183751209 | 183893656 | 142.4 | 0/3 | 2.23 | - | Unsupported deletion |
| gap_03 | NC_071442.1 | 183894153 | 183964695 | 70.5 | 0/3 | 0.892 | - | Unsupported deletion |
| gap_04 | NC_071445.1 | 0 | 447147 | 447.1 | 1/3 | 0.00399 | 17 | Putative gain in *C. jacchus* |
| gap_05 | NC_071445.1 | 30366821 | 30792573 | 425.8 | 0/2 | 0.0473 | 0 | Putative gain in *C. jacchus* |
| gap_06 | NC_071446.1 | 62986067 | 63051684 | 65.6 | 0/1 | 0.443 | - | Unsupported deletion |
| gap_07 | NC_071446.1 | 63052678 | 63073147 | 20.5 | 0/1 | 0.0048 | 0 | Putative gain in *C. jacchus* |
| gap_08 | NC_071446.1 | 63361714 | 63405120 | 43.4 | 0/1 | 0.172 | - | Unsupported deletion |
| gap_09 | NC_071447.1 | 28886324 | 29228104 | 341.8 | 0/1 | 0.00682 | 0 | Putative gain in *C. jacchus* |
| gap_10 | NC_071450.1 | 60596128 | 60783911 | 187.8 | 4/6 | 0.000215 | 2.6 | Putative gain in *C. jacchus* |
| gap_11 | NC_071450.1 | 60818423 | 60857334 | 38.9 | 3/0 | 0.00443 | 100 | Supported deletion |
| gap_12 | NC_071451.1 | 51105870 | 51126930 | 21.1 | 0/1 | 0.00497 | 100 | Supported deletion |
| gap_13 | NC_071452.1 | 110615630 | 110698577 | 82.9 | 5/0 | 0.00599 | 100 | Supported deletion |
| gap_14 | NC_071452.1 | 110719051 | 110745567 | 26.5 | 2/0 | 0.00145 | 100 | Supported deletion |
| gap_15 | NC_071452.1 | 110956057 | 111120820 | 164.8 | 12/0 | 0.000285 | 98.6 | Supported deletion |
| gap_16 | NC_071456.1 | 83463850 | 83592448 | 128.6 | 1/2 | 0.00874 | 37.4 | Putative gain in *C. jacchus* |
| gap_17 | NC_071458.1 | 0 | 126101 | 126.1 | 0/2 | 9.59 | - | Unsupported deletion |
| gap_18 | NC_071458.1 | 496643 | 1187578 | 690.9 | 0/2 | 0.198 | - | Unsupported deletion |
| gap_19 | NC_071460.1 | 34066643 | 34102832 | 36.2 | 2/0 | 0.009 | 100 | Supported deletion |
| gap_20 | NC_071460.1 | 34105452 | 34140904 | 35.5 | 2/0 | 0.00433 | 100 | Supported deletion |
| gap_21 | NC_071460.1 | 34155472 | 34266142 | 110.7 | 3/3 | 0.000433 | 100 | Supported deletion |
| gap_22 | NC_071460.1 | 34298428 | 34438616 | 140.2 | 5/2 | 0.00235 | 100 | Supported deletion |

**Supplementary Table 5. Orthology of *Callithrix jacchus* ORs within supported *Saimiri boliviensis* deletion regions.** *C. jacchus* ORs are shown with their structural status, OrthoFinder orthogroup, and inferred *S. boliviensis* orthologue(s). OR IDs are internal BITACORA-based locus identifiers; a dash indicates no inferred *S. boliviensis* orthologue.

| Candidate | *C. jacchus* OR ID | Status | *Orthogroup* | *S. boliviensis* orthologue(s) |
|---|---|---|---|---|
| gap_11 | ORsg007 | Intact | OG0000467 | — |
| gap_11 | ORsg008 | Intact | OG0000448 | — |
| gap_11 | ORsg538 | Intact | OG0000434 | — |
| gap_12 | ORsg335 | Pseudogene | OG0000333 | — |
| gap_13 | ORsg023 | Intact | OG0000321 | — |
| gap_13 | ORsg040 | Intact | OG0000452 | — |
| gap_13 | ORsg295 | Intact | OG0000581 | — |
| gap_13 | ORsg407 | Intact | OG0000324 | — |
| gap_13 | ORsg417 | Intact | OG0000310 | — |
| gap_14 | ORsg029 | Intact | OG0000397 | — |
| gap_14 | ORsg201 | Intact | OG0000392 | — |
| gap_15 | ORsg057 | Intact | OG0000381 | — |
| gap_15 | ORsg084 | Intact | OG0000575 | — |
| gap_15 | ORsg148 | Intact | OG0000460 | ORsg425 |
| gap_15 | ORsg160 | Intact | OG0000414 | — |
| gap_15 | ORsg191 | Intact | OG0000351 | — |
| gap_15 | ORsg365 | Intact | OG0000712 | — |
| gap_15 | ORsg391 | Intact | OG0000033 | — |
| gap_15 | ORsg404 | Intact | OG0000011 | ORsg414 |
| gap_15 | ORsg429 | Intact | OG0000313 | — |
| gap_15 | ORsg437 | Intact | OG0000712 | — |
| gap_15 | ORsg458 | Intact | OG0000008 | — |
| gap_15 | ORsg553 | Intact | OG0000015 | — |
| gap_19 | ORsg185 | Intact | OG0000427 | — |
| gap_19 | ORsg498 | Intact | OG0000318 | — |
| gap_20 | ORsg004 | Intact | OG0000227 | — |
| gap_20 | ORsg465 | Intact | OG0000330 | — |
| gap_21 | ORsg009 | Pseudogene | OG0000000 | ORsg394 |
| gap_21 | ORsg073 | Pseudogene | OG0000651 | — |
| gap_21 | ORsg082 | Intact | OG0000000 | ORsg394 |
| gap_21 | ORsg102 | Pseudogene | OG0000525 | — |
| gap_21 | ORsg244 | Intact | OG0000513 | — |
| gap_21 | ORsg608 | Intact | OG0000000 | ORsg394 |
| gap_22 | ORsg080 | Intact | OG0000552 | — |
| gap_22 | ORsg103 | Intact | OG0000518 | — |
| gap_22 | ORsg126 | Intact | OG0000447 | — |
| gap_22 | ORsg358 | Intact | OG0000540 | — |
| gap_22 | ORsg363 | Pseudogene | OG0000446 | — |
| gap_22 | ORsg376 | Pseudogene | OG0000359 | — |
| gap_22 | ORsg441 | Intact | OG0000595 | — |

**Supplementary Table 6. Fate of ancestral OR orthogroups across platyrrhine species with high-quality genome assemblies.** The ancestral platyrrhine OR repertoire was reconstructed using Dollo parsimony and comprised 519 orthogroups containing at least one intact OR. For each species, ancestral orthogroups were classified as retained when an intact OR was present, pseudogenized when only pseudogene copies were present, or absent when neither intact nor pseudogene copies were detected. Values are shown as number of orthogroups and percentage of the reconstructed ancestral repertoire.

| Species | Retained, n (%) | Pseudogenized, n (%) | Absent, n (%) |
|---|---|---|---|
| Aotus nancymaae | 381 (73.4) | 39 (7.5) | 99 (19.1) |
| Sapajus apella | 373 (71.9) | 59 (11.4) | 87 (16.8) |
| Cebus albifrons | 354 (68.2) | 67 (12.9) | 98 (18.9) |
| Callithrix jacchus | 346 (66.7) | 50 (9.6) | 123 (23.7) |
| Pithecia pithecia | 343 (66.1) | 88 (17.0) | 88 (17.0) |
| Plecturocebus cupreus | 320 (61.7) | 68 (13.1) | 131 (25.2) |
| Brachyteles arachnoides | 323 (62.2) | 67 (12.9) | 129 (24.9) |
| Saimiri boliviensis | 291 (56.1) | 40 (7.7) | 188 (36.2) |

**Supplementary Table 7. Supported focal OR duplication clades in *Aotus nancymaae* and capuchins.** Clades were retained at ultrafast bootstrap ≥95%. For *Aotus*, at least two ORs were required; for capuchins, at least three, given that strict orthology predicts two ORs.

| Lineage | Clade | OR composition | UFBootstrap support |
|---|---|---|---|
| Capuchins | C1 | 2 Cebus + 1 Sapajus | 100 |
| Capuchins | C2 | 2 Cebus + 1 Sapajus | 99 |
| Capuchins | C3 | 2 Cebus + 1 Sapajus | 98 |
| Capuchins | C4 | 2 Cebus + 1 Sapajus | 95 |
| Aotus | A1 | 2 Aotus | 100 |
| Aotus | A2 | 2 Aotus | 96 |

**Supplementary Table 8. PGLS models testing allometry and associations between functional OR counts and olfaction-related anatomical traits across 33 platyrrhine species.** Tested olfactory traits are olfactory bulb volume (OBV), turbinal surface area (TSA), cribiform plate area (CPA) and ethmoturbinal number (nET). Models including body mass (BM) or endocranial volume (ECV) evaluate size-adjusted associations. OR counts and continuous morphological variables were log-transformed before model fitting. β is the regression slope, p indicates model significance, $R^2$ is the adjusted R squared, and lambda is the Pagel's phylogenetic signal estimated for each model. Bold p-values indicate $p < 0.05$.

| Model | $R^2$ | λ | Predictor | slope (β) | p |
|---|---|---|---|---|---|
| log(OBV) ~ log(BM) + log(ECV) | 0.634 | 0.943 | log(BM) | 0.192 | 0.217 |
| | | | **log(ECV)** | **0.411** | **0.043** |
| log(ORs) ~ log(OBV) + log(BM) + log(ECV) | 0.520 | 0.623 | **log(OBV)** | **0.192** | **< 0.001** |
| | | | log(BM) | -0.056 | 0.085 |
| | | | log(ECV) | -0.039 | 0.327 |
| log(ORs) ~ log(OBV) + log(BM) + log(ECV) + *Saimiri* | 0.784 | 0.265 | **log(OBV)** | **0.176** | **< 0.001** |
| | | | **log(BM)** | **-0.090** | **< 0.001** |
| | | | log(ECV) | 0.005 | 0.874 |
| | | | ***Saimiri*** | **-0.126** | **< 0.001** |
| log(ORs) ~ log(OBV) + log(BM) + log(ECV) + *Aotus* | 0.533 | 0.597 | **log(OBV)** | **0.198** | **< 0.001** |
| | | | log(BM) | -0.056 | 0.087 |
| | | | log(ECV) | -0.043 | 0.336 |
| | | | *Aotus* | -0.006 | 0.906 |
| log(OBV) ~ log(ORs) + log(BM) + log(ECV) | 0.883 | 0.414 | **log(ORs)** | **2.856** | **< 0.001** |
| | | | **log(BM)** | **0.256** | **0.019** |
| | | | **log(ECV)** | **0.264** | **0.047** |
| log(OBV) ~ log(ORs) + log(BM) + log(ECV) + *Saimiri* | 0.942 | 0.000 | **log(ORs)** | **3.642** | **< 0.001** |
| | | | log(BM) | **0.358** | **0.001** |
| | | | log(ECV) | 0.149 | 0.232 |
| | | | *Saimiri* | 0.285 | 0.052 |
| log(OBV) ~ log(ORs) + log(BM) + log(ECV) + *Aotus* | 0.958 | 0.000 | **log(ORs)** | **2.572** | **< 0.001** |
| | | | **log(BM)** | **0.183** | **0.015** |
| | | | **log(ECV)** | **0.360** | **< 0.001** |
| | | | ***Aotus*** | **0.365** | **< 0.001** |
| log(ORs) ~ log(CPA) + log(BM) + log(ECV) | 0.163 | 0.913 | **log(CPA)** | **0.059** | **0.045** |
| | | | log(BM) | -0.046 | 0.252 |
| | | | log(ECV) | 0.028 | 0.568 |
| log(ORs) ~ log(TSA) + log(BM) | 0.037 | 0.937 | log(TSA) | 0.023 | 0.378 |
| | | | log(BM) | 0.020 | 0.921 |
| log(ORs) ~ nET | 0.253 | 0.879 | **nET** | **0.071** | **0.003** |
